# Dopamine functionalized, red carbon quantum dots for *in vivo* bioimaging, cancer therapeutics, and neuronal differentiation

**DOI:** 10.1101/2023.06.16.545347

**Authors:** Pankaj Yadav, Dawson Benner, Ritu Varshney, Krupa Kansara, Krupa Shah, Landon Dahle, Ashutosh Kumar, Rakesh Rawal, Sharad Gupta, Dhiraj Bhatia

## Abstract

One of the crucial requirements of quantum dots for biological applications is their surface modifications for very specific and enhanced biological recognition and uptake. Toward this, we present the green synthesis of bright, red-emitting carbon quantum dots derived from mango leaf extract (mQDs). These mQDs are conjugated electrostatically with dopamine to form mQDs-dopamine (mQDs: DOPA) bioconjugates. Bright red fluorescence of mQDs was used for bioimaging and uptake in multiple cell lines, tissues, and *in vivo* models like zebrafish. mQDs exhibited the highest uptake in brain tissue as compared to others. mQD:DOPA conjugate induced cellular toxicity only in cancer cells while showing increased uptake in epithelial cells and zebrafish. Additionally, the mQDs: DOPA promoted neuronal differentiation of SH-SY5Y cells to complete neurons. Both mQDs and mQDs: DOPA exhibited potential for higher collective cell migrations implicating their future potential as next-generation tools for advanced biological and biomedical applications.

**TOC:** mQDs were electrostatically conjugated with dopamine (DOPA) to form the mQDs: DOPA bioconjugate. mQDs are used to image cells, tissues, and zebrafish embryos. mQDs: DOPA kills cancer cells, differentiates neuronal cells, and increases the uptake of mQDs in zebrafish embryos.

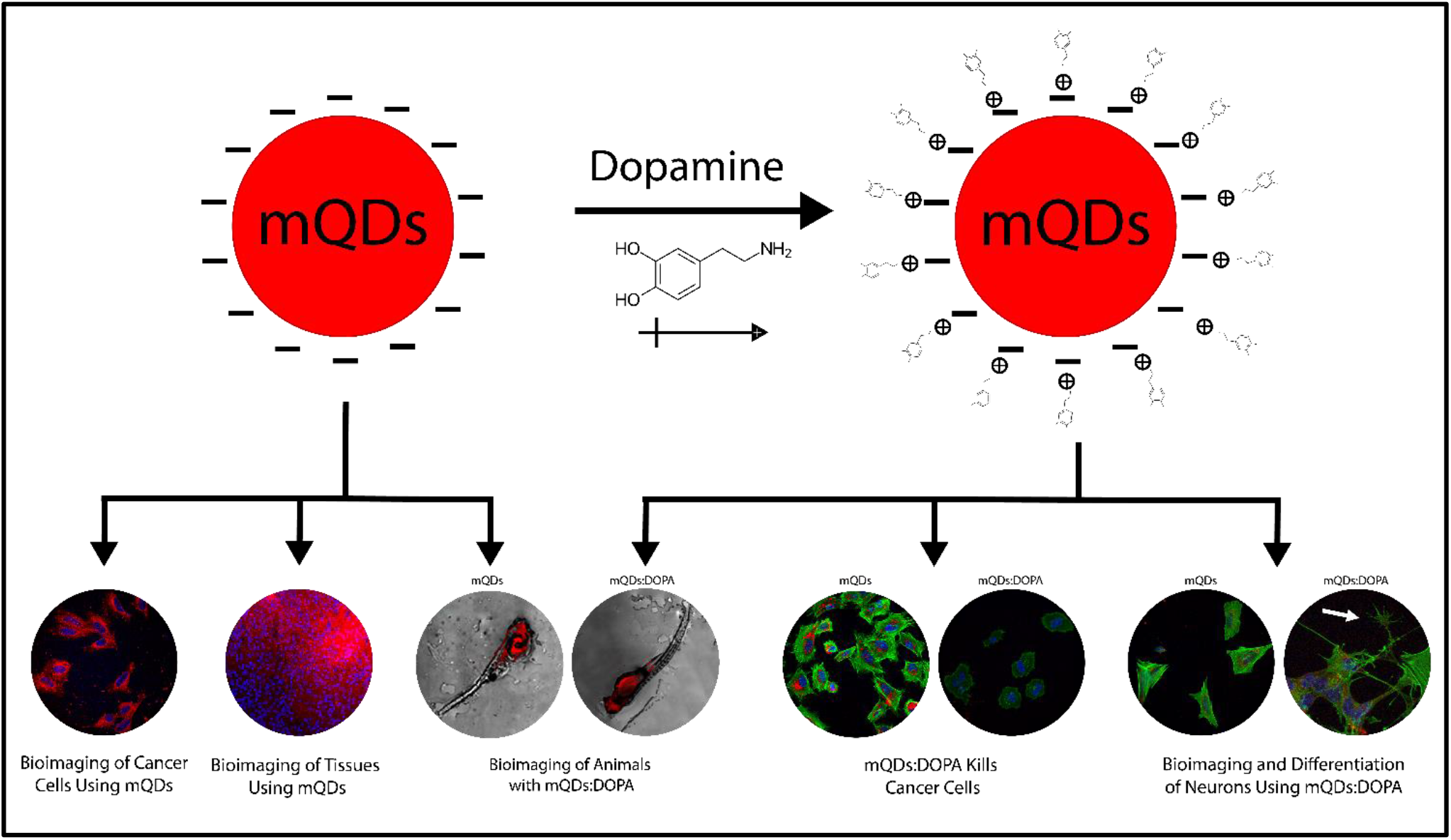

## Introduction

Neuronal differentiation is the first step of neuron development in which cells produce more dendrites and longer axons to connect with other nerve cells. This process plays a significant role in life, and damaged neurons can contribute to Alzheimer’s and Parkinson’s disease, among others1,2. Different neurotransmitters, including the thyroid hormine^3^, SOX transcription factors^4^, retinoic acid, fibroblast growth factors^5,^ and dopamine, have been identified as critical for differentiation. Among these, dopamine’s effect in neuronal differentiation remains the least well understood; while only limited regions of the brain produce it, dopamine is required for most elements of the nervous system^1,6^.

Quantum dots (QDs) are zero-dimensional particles, typically below 10 nanometers in size. Surface modifications imbue them with peculiar qualities like high photon yield, controlled electron transport controlling their fluorescence properties, and increased uptake in biological systems^7^. QDs synthesized from CdSe/ZnS have been conjugated with dopamine and subsequently used for intercellular and in vitro pH sensing^8^. Polydopamine-coated gold nanoparticles interact with the D2DR dopamine receptor through amine and catechol functional group^9^. Dopamine mediates the cell-nanoparticle interactions and, therefore, can be used as a component for targeted nanoscale drug delivery. However, all of these nanomaterials are inorganic or metallic thereby posing challenges to their applications in biological systems. To our knowledge, there are no current studies involving dopamine binding to carbon based organic quantum dots and their subsequent potential in neuronal differentiation.

In this study, we have synthesized a new class of multipurpose red-emitting carbon dots derived from mango leaves (mQDs) with an emission wavelength of 670 nm and size of 2-5 nm. The mQDs were used for bioimaging of cancer cells (HeLa and SUM159A cells), retinal pigment epithelial cells (RPE-1 cells), and neuronal cells (SH-SY5Y), both differentiated and undifferentiated. mQDs were used for imaging tissue specific uptake from mouse brain, heart, liver, and kidney, as well as for *in vivo* model systems such as zebrafish embryos. The synthesized mQDs were capable of surface modification by dopamine (DOPA) through electrostatic interaction to form a bioconjugate (mQDs: DOPA). The bioconjugate exhibited specific toxicity towards cancer cells (SUM 159A) and enhanced uptake in epithelial cells and zebrafish. Further, the conjugate promoted neuronal cell differentiation of SH-SY5Y cells. Finally, both mQDs and mQDs: DOPA positively affected cell migration in RPE-1 cells. These new class of quantum dots with excellent fluorescent properties, robust surface chemistry for modifications can emerge as new class of bioimaging and theranostic tools for biological and biomedical applications.

### Synthesis, Characterization, and Conjugation of mQDs

Fluorescent red mango quantum dots (mQDs) were synthesized from mango leaves. Fresh leaves were plucked and then dried at room temperature. The dried mango leaves were then powdered using a grinder. The obtained leaf powder was dissolved in ethanol (in a w/v ratio of 1:10) and stirred for 4 hours at room temperature. The solution was centrifuged at 10,000 rpm at room temperature to obtain the mango leaf extract, discarding the sediment. The mango leaf extract was refluxed for 2 hours at 160°C to form mQDs. The obtained mQDs solution was then syringe filtered using a 0.22 μm filter. The mQDs appeared green in room light and red under UV light (**Supplementary Figure S1**). The ethanol was evaporated using a rotavapor to obtain a powder of mQDs, characterized using various techniques to identify their optical and surface properties.

The absorbance spectra of mQDs showed distinct peaks at 279nm, 369nm, and 667nm, as shown in **Figure 1A**. The peak at 279m corresponds to the n-π* electronic transition in carbon structure, and 365nm corresponds to the surface or molecule center of the mQDs. The peak at 667 nm is attributed to the chlorophyll present in green plants. The emission spectra revealed the excitation-dependent emission of mQDs. The fluorescence intensity increased with starting excitation from 300nm, with maxima at 400nm and subsequent decrease, as shown in **Figure 1B**. The FTIR spectra showed the presence of the following surface groups on mQDs: 3219 cm^-1^ for alcohol-OH; 2925 cm^-1^ and 2851 cm^-1^ for sp3 –CH stretch; 1707 cm^-1^ for carbonyl -C=O; 1609 cm^-1^ for alkene C=C; 1451 cm^-1^ for sp3–CH bend; 1323 cm^-1^; 1200 cm^-1^ for ester C-O; 1032 cm^-1^ for alcohol C-O; and 880 cm^-1^, 808 cm^-1^ and 763 cm^-1^ for sp2 –CH bend (**Figure 1C**). The comparison of mango leaves peaks at 3284 cm^-1^ (alcohol-OH), 2919 cm^-1^ (sp3 –CH stretch), 2849 cm^-1^ (sp3 –CH stretch), 1612 cm^-1^ (alkene C=C), and 1033 cm^-1^ (alcohol C-O) with mQDs shows the shift of peaks in mQDs at 3219 cm^-1^, 2925 cm^-1^, 2851 cm^-1^, 1609 cm^-1^, 1032 cm^-1^. This comparison also reveals the formation of new bonds in mQDs at 1707 cm^-1^, 1451 cm^-1^, 1323 cm^-1^, 1200 cm^-1^, and 880 cm^-1^. The quasi-spherical morphology and topography were studied with the help of atomic force microscopy (AFM) in **Figure 1D**. AFM analysis of mQDs showed their size and height at around 10 nm and 2 nm, respectively (**Figure 1E, F)**. The 3-dimensional view and height profile of the mQDs is shown in **Figure 1F, G**, and **Supplementary Figure S2**. The powder XRD analysis showed the amorphous nature of mQDs with broad peaks at 18° (**Figure 1H)**. Dynamic light spectroscopy revealed that mQDs have a diameter of 2.5 nm in ethanol as solvent. The quantum yield of the mQDs was calculated to be 202% of Rhodamine B’s quantum yield, as shown in **Supplementary Figure S3A, B**. The maximum excitation spectra of mQDs are at 410 nm, which overlaps with the absorption spectra (**Supplementary Figure S3C, D**). The lifetime decay of electrons in the excited state was 20 ns; this shows the radiative recombination mechanism of luminescence^10^, in **Supplementary Figure S3D**. The charge on the mQDs was measured to be –32.4 mV in water in **Supplementary Figure S4D**.

**Figure 1:**
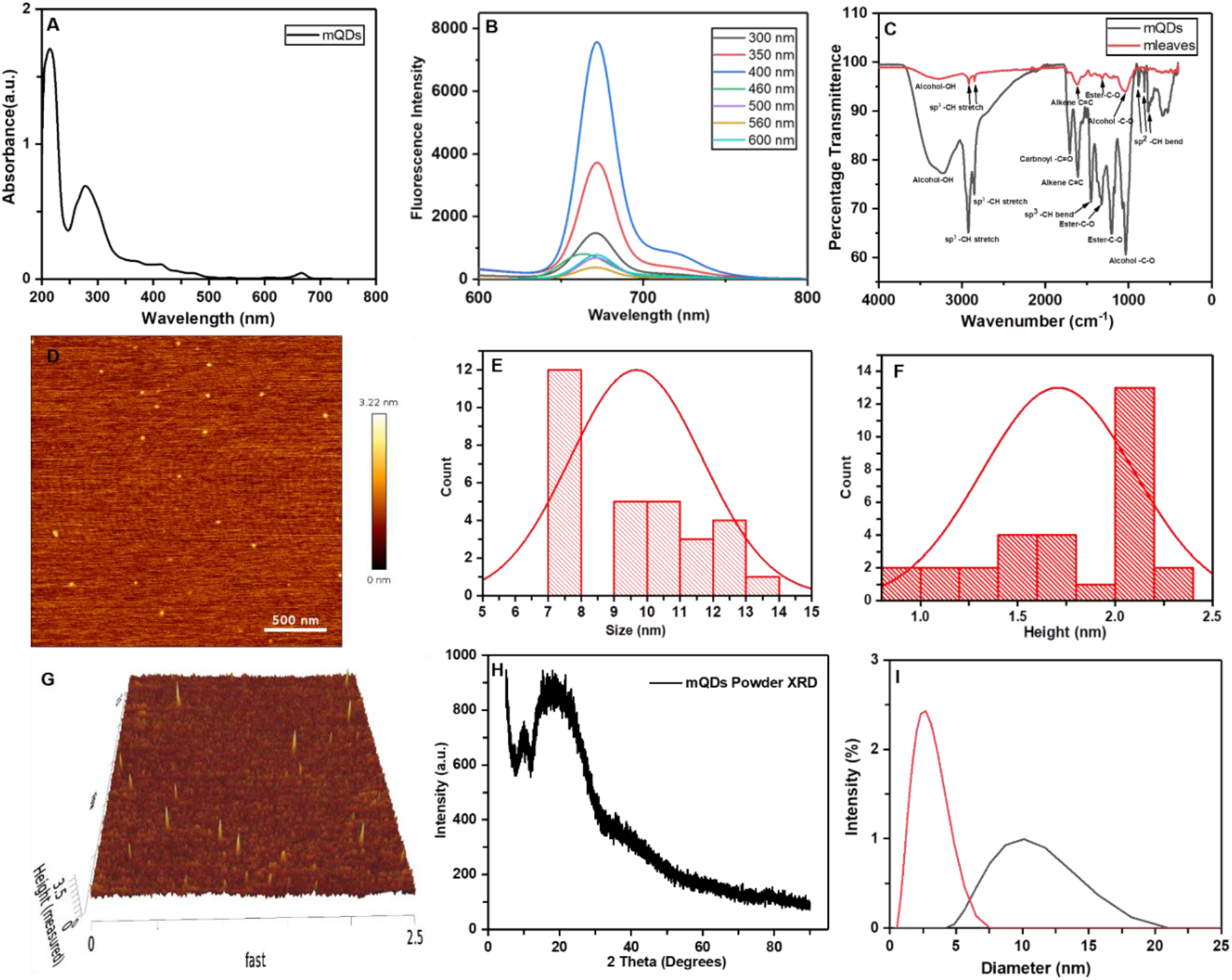
Characterization of mQDs. **(A)** The absorbance spectra of mQDs showed distinct peaks at 279nm, 369nm, and 667nm. **(B)** mQDs showed excitation-dependent emission at 670nm; fluorescence intensity increases achieved a maximum at 400 nm and subsequent decrease **(C)** The FTIR spectra indicate the presence of surface groups on mQDs: 3219 cm^-1^ for alcohol-OH; 2925 cm^-1^ and 2851 cm^-1^ for sp^3^ –CH stretch; 1707 cm^-1^ for carbonyl -C=O; 1609 cm^-1^ for alkene C=C; 1451 cm^-1^ for sp^3^ –CH bend; 1323 cm^-1^ and 1200 cm^-1^ for ester C-O; 1032 cm^-1^ for alcohol C-O; and 880 cm^-1^, 808 cm^-1^, and 763 cm^-1^ for sp^2^ –CH bend. The comparison of mango leaves peaks at 3284 cm^-1^ (alcohol-OH); 2919 cm^-1^ (sp^3^ –CH stretch); 2849 cm^-1^ (sp^3^ –CH stretch); 1612 cm^-1^ (alkene C=C); and 1033 cm^-1^ (alcohol C-O) with mQDs shows the shift of peaks in mQDs at 3219 cm^-1^, 2925 cm^-1^, 2851 cm^-1^, 1609 cm^-1^, and 1032 cm^-1^, along with the formation of new bonds in mQDs at 1707 cm^-1^, 1451 cm^-1^, 1323 cm^-1^, 1200 cm^-1^, and 880 cm^-1^. **(D)** The quasi-spherical morphology and topography of mQDs were studied using atomic force microscopy (AFM). **(E)** Histogram plot of AFM analysis of mQDs showing their size at around 10 nm; n=30. **(F)** Histogram plot of AFM analysis of mQDs showing their height at about 2 nm; n = 30. **(G)** The 3-dimensional view of the mQDs. **(H)** The powder XRD analysis shows the amorphous nature of mQDs with broad peaks at 18°. **(I)** DLS analysis of mQDs in ethanol as solvent shows the hydrodynamic radius of mQDs as 2.5 nm.

The negative charge on the mQDs prompted us to conjugate electrostatically with the positively charged small molecule dopamine (DOPA). The mQDs and DOPA were mixed in 1:2, 1:5, 1:10, and 1:20 ratios with water as the solvent. The hydrodynamic size and zeta potential were determined by the conjugation of mQDs with DOPA (**Supplementary Figure S4D**). An absorbance peak at 280 nm indicated successful conjugation of mQDs and DOPA (**Supplementary Figure S4C**). The size of only mQDs and DOPA were 37.8 nm and 0.719 nm, respectively (**Figure 2A, B**). The size of the conjugate in 1:2, 1:5, 1:10, and 1:20 ratios were 43.8 nm, 0.621 nm, 0.719 nm, and 0.833 nm, respectively **Figure 2C, D, E, and F**. The conjugate in ratios 1:5, 1:10, and 1:20 showed the size corresponding to that of DOPA alone, showing that dopamine masked the mQDs. However, in the case of the ratio 1:2, the hydrodynamic size increased to 43.8 nm, indicating the association of DOPA with mQDs and thus formation of a zwitterionic molecule. Therefore, the 1:2 ratio of mQDs and DOPA was utilized; from here on, the 1:2 ratio is referred to as mQDs: DOPA unless otherwise stated. The AFM analysis of mQDs: DOPA revealed a shell-like structure of dopamine which circled the mQDs, confirming the mQDs: DOPA conjugation as shown in **Figure 2G, H**. FTIR analysis of mQDs: DOPA showed the disappearance of the peaks at 1707 cm^-1^ (carbonyl -C=O); 1451 cm^-1^ (sp3 –CH bend); 1323 cm^-1^; (ester C-O) and 880 cm^-1^, 808 cm^-1^, 763 cm^-1^ (sp^2^ –CH bend) compared to mQDs. In addition, there is a shift in the percentage transmittance; mQDs: DOPA has stronger transmittance than mQDs alone but weaker transmittance than DOPA alone, suggesting the mQDs: DOPA conjugation. DOPA loading onto mQDs was subsequently calculated to be 52.47% (**Supplementary Figure S4**).

**Figure 2:**
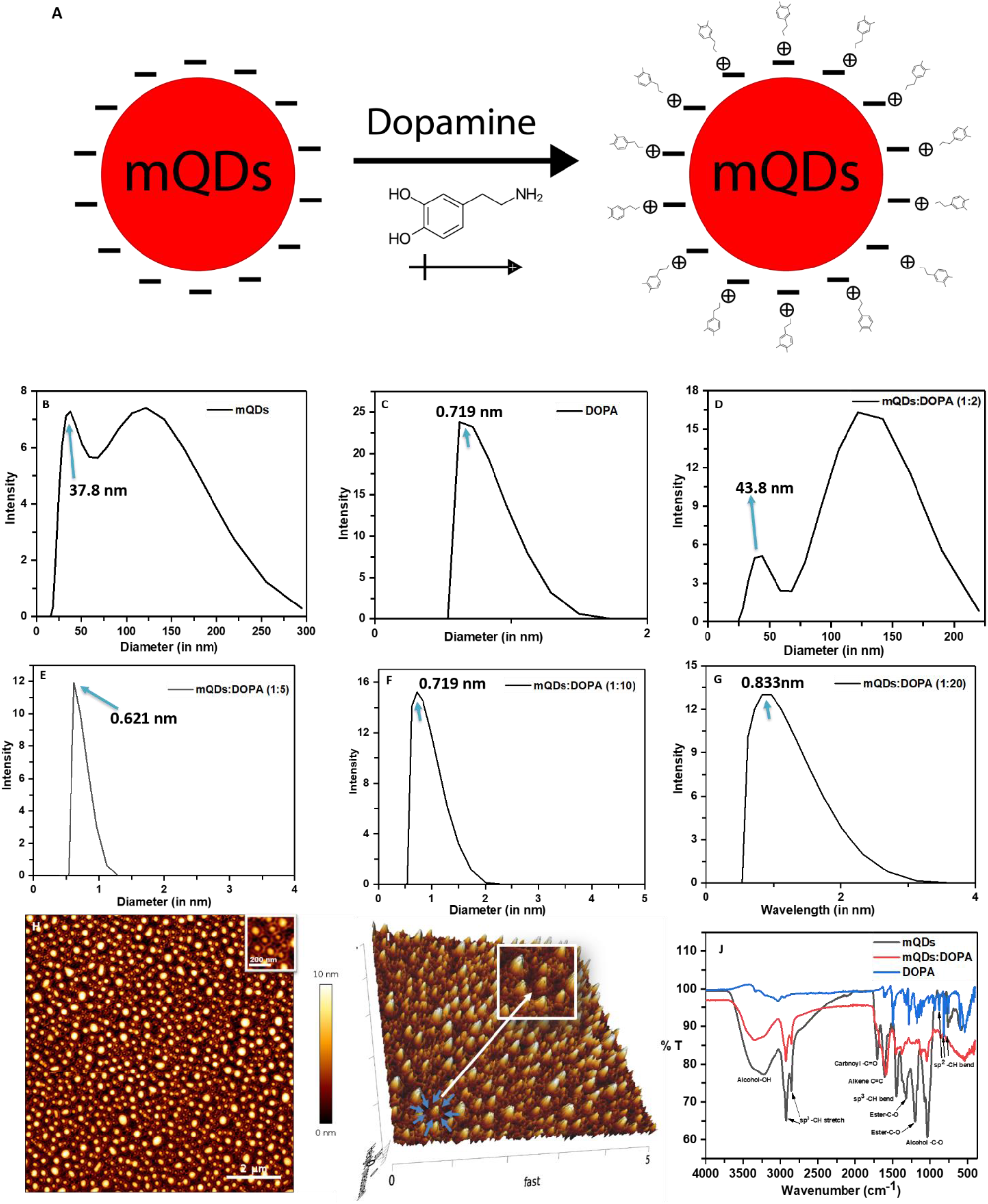
Conjugation of mQDs with dopamine (DOPA) **(A)** Scheme of conjugation of mQDs (-) with DOPA (+) through electrostatic interaction. **(B)** The hydrodynamic radius of mQDs alone in water as solvent was measured as 37.8 nm. **(C)** The hydrodynamic radius of DOPA alone in water as solvent was measured as 0.719 nm. **(D)** The hydrodynamic radius of mQDs: DOPA (1:2) in water as solvent was measured as 43.8 nm. **(E)** Hydrodynamic radius of mQDs: DOPA (1:5) in water as solvent measured as 0.621 nm. **(F)** Hydrodynamic radius of mQDs: DOPA (1:10) in water as solvent measured 0.719 nm. **(G)** Hydrodynamic radius of mQDs: DOPA (1:20) in water as solvent measured as 0.833 nm. **(H)** The AFM analysis of mQDs: DOPA forming a cage-like structure with dopamine sitting at the periphery of mQDs, confirming the conjugation of mQDs with DOPA. **(I)** 3-Dimensional view of the DOPA circling mQDs, indicated with the arrows.

### mQDs uptake in mammalian cells, tissues, and zebrafish

To study the cell-specific behavior of mQDs and mQDs: DOPA, we performed the 3-[4,5-dimethylthiazol-2-yl]-2,5-diphenyltetrazolium bromide (MTT) assay on epithelial cells (RPE1), breast cancer cells (SUM159A) and neuronal cells (SH-SY5Y)^11^. An increasing concentration of mQDs (100, 200, 300, 400, 500 μg/mL) and mQDs: DOPA (100, 200, 300, 400, 500 μg/mL) was added to the RPE1 cells, SUM159A cells, and SH-SY5Y cells. The percentage viability was calculated after 24 hours of incubation.

The cell viability of RPE1 incubated with mQDs alone increased from 93.22% in 100 μg/mL to 108.05 % in 500 μg/mL, suggesting the proliferating nature of mQDs in the case of non-cancerous cells (**Supplementary Figure S5A**). This could also indicate the use for wound-healing properties of mQDs. In the case of SUM159A cells, the viability of cells varied with dosage. 100, 200, 300, 400, and 500 μg/mL yielded 49.21%, 46.65%, 50.53%, 41.84%, and 47.54% cell viability (**Supplementary Figure S5C**). The observation in cancer cells shows that mQDs have anti-cancer properties, coinciding with previous studies on the anti-cancer properties of mango leaves. The percentage cell viability of mQDs in SHSY-5Y cells was 80.41%, 129.37%, 91.6%, 86.71%, and 109.09% for 100, 200, 300, 400, and 500 μg/mL, respectively (**Supplementary Figure S5E**). These indicate suitability and safety in non-cancerous cells. The cell viability of RPE1 cells incubated with mQDs: DOPA showed cell viability of 103.91%, 134.80%, 133.93%, 112.55%, and 154.049% for 100, 200, 300, 400, and 500 μg/mL respectively (**Supplementary Figure S5B**). In the case of SUM159A cells, the cell viability was 48.50%, 55.52%, 50.66%, 48.26%, and 47.01% for 100, 200, 300, 400, and 500 μg/mL of mQDs: DOPA respectively (**Supplementary Figure S5D**). The cell viability of SH-SY5Y cells incubated with mQDs: DOPA showed cell viability of 95.1%, 95.8%, 80.41%, 70.62%, and 104.89% for 100, 200, 300, 400, and 500 μg/mL, respectively (**Supplementary Figure S5F**). In the case of RPE1 cells, SHSY-5Y, and SUM159A, the cell viability was, respectively, 86.57%, 129.37%, and 46.65% at 200 μg/mL of mQDs and 134.80%, 95.8%, and 55.52% at 200 μg/mL of mQDs: DOPA (**Supplementary Figure S5**). The overall cell viability assay suggests the nontoxicity of mQDs and mQDs: DOPA at 200 μg/mL concentration in RPE1 and SH-SY5Y cells and a toxic effect in SUM 159A cells. Therefore, both mQDs and mQDs: DOPA have anti-cancer properties.

The red fluorescence of mQDs enabled the high-contrast imaging of different types of cells (cancer, epithelial and neuronal), tissues (brain, heart, kidney, and liver), and zebrafish embryos. The cells, tissues, and zebrafish embryos were treated with mQDs in order of increasing concentration of 50 μg/mL, 100 μg/mL, and 200 μg/mL, as shown in **Supplementary Figures S6, S7, S8, and S9**. The fluorescence intensity was quantified in ImageJ to measure the uptake of mQDs. Cancer cells (SUM159A) were treated with an increasing concentration of mQDs for 15 minutes. The mQDs were subsequently endocytosed and uptaken into cells. Increasing the concentration of mQDs resulted in increased fluorescence intensity and hence the increased uptake of mQDs, as shown in **Figure 3**. The real-time endocytosis of mQDs was observed by performing live cell imaging on GFP-labeled microtubules in HeLa cells (MT GFP HeLa). The MT GFP HeLa cells were chosen to have red fluorescent mQDs in contrast to the green fluorescence of GFP. The MT GFP HeLa cells were treated with 500 μg/mL of mQDs at t=0 minutes. As time passed, the intensity of the red fluorescence increased inside the cell and on the boundary of the MT GFP HeLa cells. The interaction of mQDs with HeLa cells can be seen at the periphery of the cell membrane, shown by arrows at times t=2, 4, and 6 minutes in **Figure 3A-D**. The results show that mQDs and mQDs: DOPA were endocytosed in increased volumes in cancerous cells, enabling accurate imaging and reduced cancer proliferation (**Figure 3E-H**).

**Figure 3:**
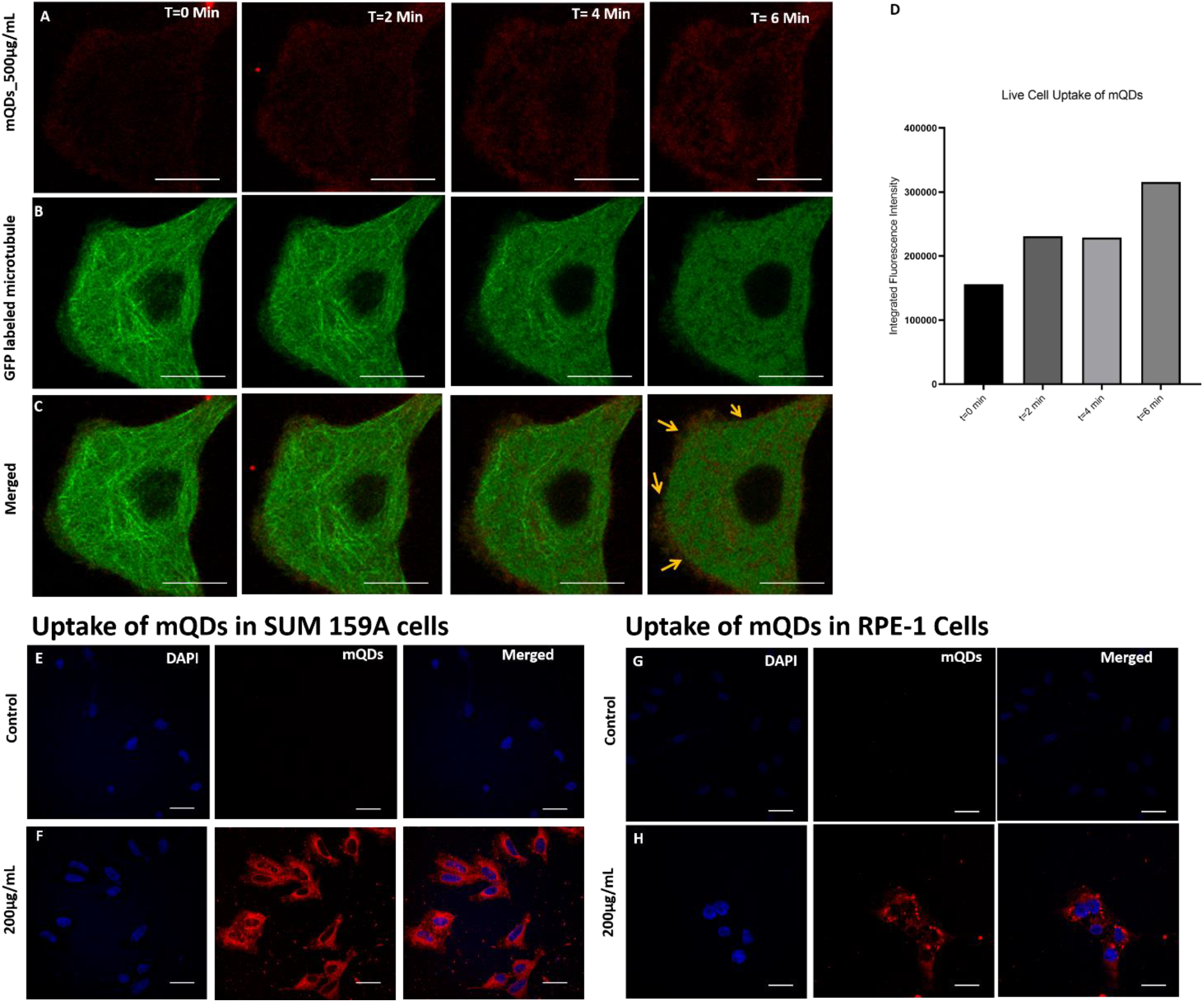
SUM 159A Cells. (scale bar 5μm), **RPE-1 cells** (scale bar 5μm) **(A)** mQDs, 500μg/mL, imaged at t=0, 2, 4, and 6 minutes. **(B) GFP-labeled** microtubule HeLa cells were imaged at t=0, 2, 4, and 6 minutes. **(C)** Merged imaging of green HeLa cells and red mQDs imaged at t=0, 2, 4, and 6 minutes. **(D)** Quantified mQDs fluorescence at t=0, 2, 4, and 6 minutes. **(E)** Control SUM159A cell imaging split into blue (DAPI labeling nucleus) and red (mQDs fluorescence) channels. **(F)** SUM159A cell imaging of 200 μg/mL mQDs dose, blue (DAPI labeling nucleus), and red (mQDs fluorescence) channels. **(G)** Control RPE-1 cell imaging, blue (DAPI labeling nucleus) and red (mQDs fluorescence) channels. **(H)** RPE-1 cell imaging of 200 μg/mL mQDs, blue (DAPI labeling nucleus), and red (mQDs fluorescence) channels.

One major issue with contemporary QDs is their inability to provide deep tissue imaging, as blue/green QDs are speculated to cause autofluorescence of samples^12^. However, red CDs easily circumvent this issue, as shown in Figures **4A and B in mouse brain tissue**. The red fluorescence of mQDs was also used for deep-tissue imaging of mouse heart, kidney, and liver tissue sections—**supplementary Figure S6**,**7**,**8**. The tissue sections were treated with mQDs for 15 minutes with an increasing concentration of 50 μg/mL, 100 μg/mL, and 200 μg/mL. The fluorescence intensity increased with increasing concentration, suggesting an increased uptake of the mQDs. Fold change is a measure used to determine uptake relative to a control value; a two-fold increase means the uptake was double the control. Upon measuring fold change, we could see that mQDs were endocytosed the most in brain tissues, followed by kidney tissues, with heart and liver tissues sharing equal fold change, as shown in **Figure 4C**. Similarly, zebrafish embryos treated with mQDs and mQDs-Dopa were imaged in **Figure 4E** and **Supplementary Figure S9**. Zebrafish embryos were treated with the same dose of mQDs and mQDs: DOPA, and the red fluorescence was quantified by measuring the fluorescence intensity in **Figure 4D**. Compared with mQDs, mQDs: DOPA showed a significant change in endocytosis. This suggests that dopamine conjugation increases the uptake of mQDs, further aiding the bioimaging *in vivo* model systems (**Figure 4**).

**Figure 4.**
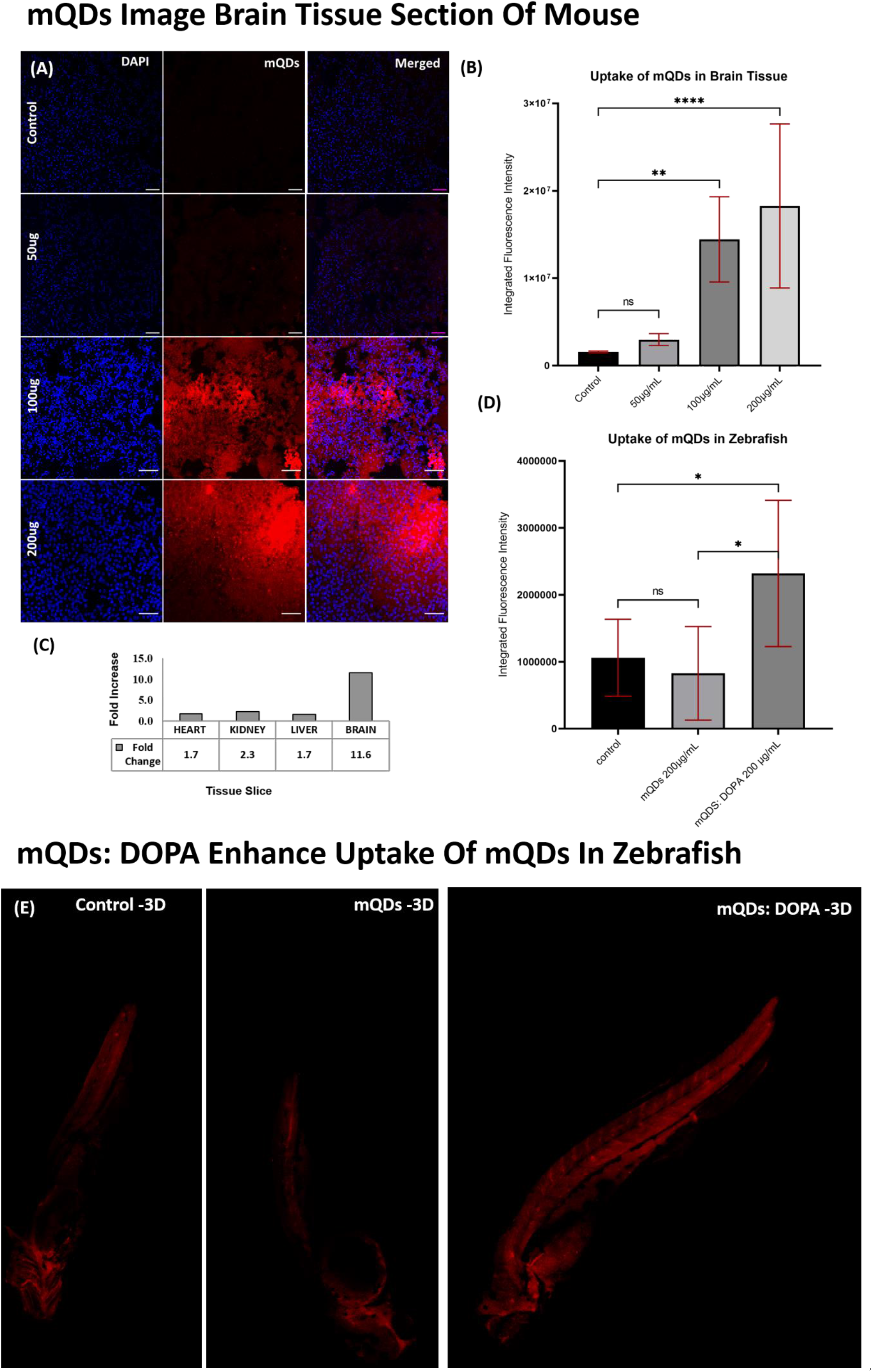
Brain tissue and Zebrafish imaging (Scale bar 5μm). **(A)** Brain tissue imaging, at concentrations varying from 0 to 200 μg/mL of mQDs, split into blue (DAPI labeling nucleus) and red (mQDs fluorescence) channels. **(B)** Quantified mQDs uptake in brain tissue. **(C)** Fold increase of red fluorescence in different tissue types. **(D)** Quantified fluorescence in zebrafish with control, mQDs, and mQDs: DOPA. **(E)** Imaging zebrafish with control, mQDs, and mQDs: DOPA. The following thresholds were used when determining significance. ***: p< 0.001. **: p< 0.01. ns: p>0.05 and denotes no significance. (n = 5 tissues sections and zebrafish embryos for quantification)

### mQDs:DOPA kills specifically cancer cells

Because of widely reported anti-cancer properties in CDs ^13,14^, the effect of mQDs: DOPA was studied on cancerous cell types. Fluorescence of mQDs and standard markers (phalloidin for actin and DAPI for nucleus) helped to visualize the cellular morphology and physiology. Based on the toxicity studies, we designed an experiment to validate our results. Three types of cells were taken for the study, namely cancer cells (SUM159A), epithelial cells (RPE-1), and neuronal cells (SHSY-5Y). The cytoskeleton structure of the cells was labeled phalloidin; the nucleus was stained with DAPI, and red fluorescence confirmed the uptake of both mQDs and mQDs: DOPA. The cells were seeded and treated with mQDs and mQDs: DOPA to observe the effect.

In the case of cancer cells, there was an increase in uptake of 200 μg/mL of mQDs compared to 100 μg/mL and control, as shown in **Figures 5A, B, C**. However, when cancer cells were treated with mQDs: DOPA, the cytoskeleton structure of the cells was ruptured (phalloidin channel), and mQDs were not endocytosed in the cells (red channel), as shown in **Figure 5D**. The measured fluorescence intensity of both phalloidin and mQDs shows a significant decrease (**Figure 5E and F**) and rupturing of cytoskeleton structure in the case of mQDs: DOPA. In the case of epithelial cells, there was an increase in uptake of 200 μg/mL of mQDs compared to 100 μg/mL and control, as shown in **Figures 5G, H1, and I**. However, when epithelial cells were treated with mQDs: DOPA, the cytoskeleton structure of the cells was not ruptured (phalloidin channel); moreover, the endocytosis of mQDs increased in the cells (red channel), as shown in **Figure 5J**. The fluorescence intensity of phalloidin shows a maintained cytoskeleton structure and a significant increase of mQDs in the case of mQDs: DOPA (**Figures 5K and L**). In the case of neuronal cells, there was an increase in uptake of 200 μg/mL of mQDs compared to 100 μg/mL, 50 μg/mL, and control, as shown in **Supplementary Figure S10E**. When neuronal cells were treated with mQDs: DOPA, the endocytosis of mQDs in the cells increased (red channel), as shown in **Supplementary Figure S10**.

**Figure 5:**
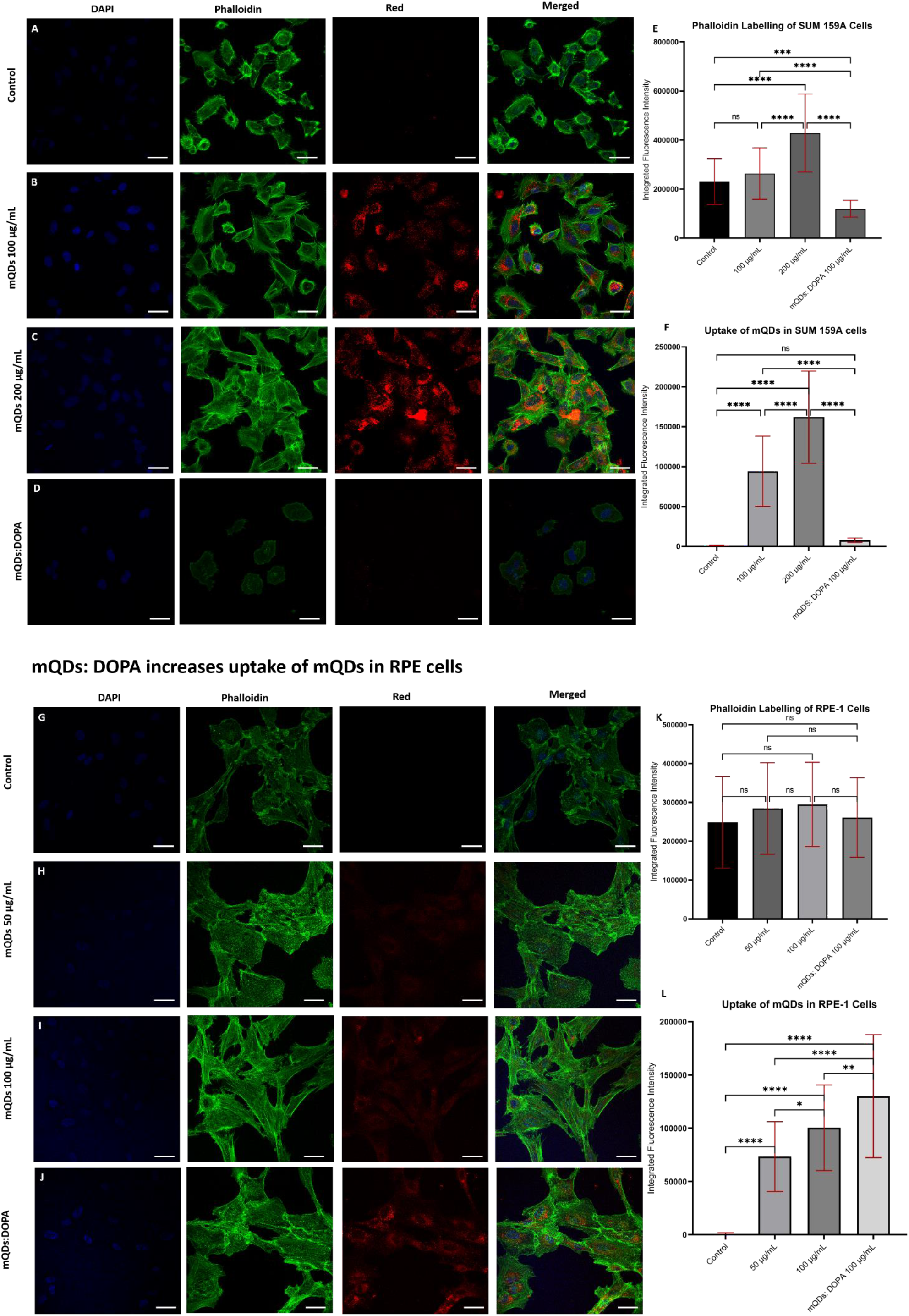
mQDs: DOPA kills cancer (SUM 159A) cells and enhances uptake of mQDs in epithelial (RPE-1) cells: DAPI labels the nucleus, phalloidin labels the actin, and red is the fluorescence of mQDs. **(A)** SUM 159A cells not treated with mQDs and mQDs: DOPA. (B) SUM 159A cells treated with 100 μg/mL of mQDs (C) SUM 159A cells treated with 200 μg/mL of mQDs. **(D)** SUM 159A cells treated with 100 μg/mL mQDs: DOPA (1:2) show a decrease in phalloidin and red fluorescence with rupturing of cytoskeleton structure. **(E)** Phalloidin labeling of SUM 159A cells, there is a significant decrease in fluorescence intensity of 100 μg/mL of mQDs: DOPA when compared with control, 100 μg/mL, 200 μg/mL of mQDs **(F)** The uptake of mQDs in SUM 159A cells increases with increasing concentration of mQDs from 100 μg/mL to 200 μg/mL and decreases significantly in case of mQDs: DOPA similar to that of control, showing the killing of cancer cells. **(G)** RPE-1 cells not treated with mQDs and mQDs: DOPA. **(H)** RPE-1 cells treated with 50 μg/mL of mQDs **(I)** SUM 159A cells treated with 100 μg/mL of mQDs. **(J)** SUM 159A cells treated with 100 μg/mL mQDs: DOPA (1:2) shows the phalloidin labeling of actin and red fluorescence without rupturing of cytoskeleton structure. **(K)** Phalloidin labeling of SUM 159A cells, there is a significant decrease in fluorescence intensity of 100 μg/mL of mQDs: DOPA when compared with control, 100 μg/mL, 200 μg/mL of mQDs **(L)** The uptake of mQDs in SUM 159A cells increases with increasing concentration of mQDs from 100 μg/mL to 200 μg/mL and decreases significantly in case of mQDs: DOPA similar to that of control, showing the killing of cancer cells—scale bar 5μm. The following thresholds were used when determining significance. ****: P≤ 0.0001. ***: p≤ 0.001. **: p≤ 0.01. *: p≤ 0.05. ns: p>0.05 and denotes no significance. (n = 50 cells for quantification)

### MQDs:DOPA promotes neuronal cell differentiation and collective cell migration

The neuronal differentiation capacity of mQD:DOPA was studied compared to control, mQDs, and DOPA. Retinoic acid (RA) was chosen as a standard to compare and validate the experiment due to its influence on neuron differentiation and development^15,16^. To check the differentiation property of mQDs: DOPA, SH-SY5Y neuronal cells were treated with 100 μg/mL of mQDs: DOPA, 100 μg/mL of mQDs, and 50 μg/mL of DOPA. The observed results in bright field and fluorescence imaging, as shown in **Figure 6A** and **Supplementary Figure S12**, show that mQDs: DOPA promote differentiation of neuronal SH-SY5Y cells. The differentiation of the cells started to be visible with outward dendrite projection from day six onwards. The treatment on day nine accelerated the cells’ differentiation **(supplementary methods section 1.9)**. It fully developed into a neuronal cell on day ten, as shown in **Figures 6A, E (E1, E2, E3), I, J, K, and M**. However, the mQDs did not exhibit quantitative differentiation, as shown in **Figures 6B, F, and 6N**. The cells were dead with DOPA treatment after day one, as shown in **Figures 6C, G, and O**. Similarly, in the case of RA, the cells started to differentiate from day two onwards and fully developed a connected network by the end of day ten. RA was used as a positive control to validate the differentiation in neuronal cells compared to mQDs: DOPA, as shown in **Figures 6D, H, and P**. The length of dendrite was quantified and observed to be 2.9395 μm, 4.5624 μm, 2.3730 μm and 4.0794 μm in control, mQDs: DOPA, mQDs, and RA, respectively. Thus, there is a significant increase in dendrite length when treated with mQDs: DOPA compared to control and mQDs, as shown in **Figure 6Q**. However, the mQDs’ uptake was higher when compared to mQDs: DOPA, as shown in **Figure 6R**.

**Figure 6:**
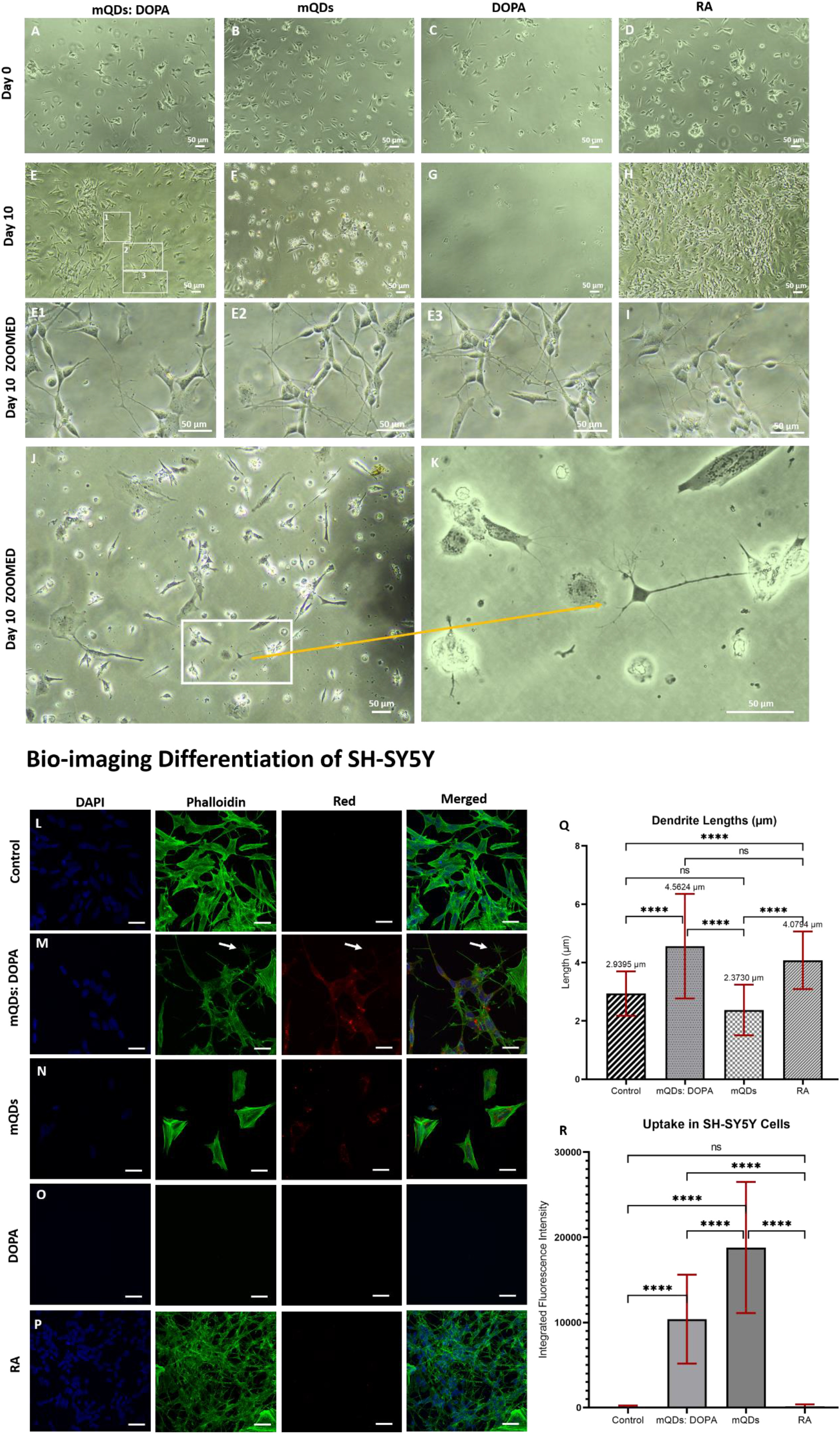
mQDs: DOPA promotes differentiation of SH-SY5Y neuroblastoma cells. **(A)** SH-SY5Y cells on day zero before the treatment with mQDs: DOPA. **(B)** SH-SY5Y cells on day zero before the treatment with mQDs. **(C)** SH-SY5Y cells on day zero before the treatment with DOPA. **(D)** SH-SY5Y cells on day zero before the treatment with RA. **(E)** SH-SY5Y cells on day ten after the treatment with mQDs: DOPA. **(F)** SH-SY5Y cells on day ten after the treatment with mQDs. **(G)** SH-SY5Y cells on day ten after the treatment with DOPA. **(H)** SH-SY5Y cells on day ten after the treatment with RA. **(E1)** Highlights region 1 of SH-SY5Y cells on day ten after the treatment with mQDs: DOPA. **(E2)** Highlights region 2 of SH-SY5Y cells on day ten after the treatment with mQDs: DOPA **(E3)** Highlights region 3 of SH-SY5Y cells on day ten after the treatment with mQDs: DOPA **(I)** SH-SY5Y cells on day ten after the treatment with mQDs: DOPA **(J)** SH-SY5Y cells on day ten after the treatment with mQDs: DOPA **(K)** Zoomed view of the rectangle region in figure J of SH-SY5Y cells on day ten after the treatment with mQDs: DOPA. **Bio-imaging Differentiation of SH-SY5Y**. Images A-H and J are taken in 10x objective. Images E1-E3, I, and K are taken at 40x objective. **(L)** The untreated cells after day ten. **(M)** mQDs: DOPA-treated cells after day ten shows the differentiated neurons pointed by the arrow. **(N)** mQDs treated cells after day ten shows their uptake but not differentiation. **(O)** DOPA-treated cells after day ten show the killing of cells. **(P)** RA-treated cells after day ten show differentiation of cells. **(Q)** The dendrite length of the neurons was measured, and the average length was 2.9395 μm, 4.5624 μm, 2.3730 μm, and 4.0794 μm in control, mQDs: DOPA, mQDs, and RA respectively. There is a significant difference in dendrite length of mQDs: DOPA compared to control and mQDs but not with RA. **(R)** The uptake of mQDs: DOPA and mQDs in differentiated cells. The uptake of mQDs is more than that of mQDs: DOPA. The scale bar is 5 μm. Images are taken in 63x oil objective. DAPI labeled the nucleus, phalloidin labeled the actin filament, and the red channel showed mQDs fluorescence of SH-SY5Y cells. The following thresholds were used when determining significance. ****: p≤ 0.0001. ***: p≤ 0.001. ns: p>0.05 and denotes no significance. (n = 50 cells for quantification)

The effect of DOPA on the uptake of mQDs in differentiated SH-SY5Y cells was studied by differentiating cells with RA (5 mM, **Supplementary Table S1**)^17^, and then treated with 100 μg/mL of mQDs: DOPA, 100 μg/mL of mQDs, and 50 μg/mL of DOPA. They were monitored over two weeks, and confocal microscopy images were taken on days 0, 7, and 14 (**Figure 7A-D, Supplementary Figure S13**). Uptake of both mQDs and mQDs: DOPA increased as time progressed (**Supplementary Figure S11**). In addition, as shown in **Figure 7E**, mQDs: DOPA showed increased uptake compared to mQDs alone by day 14. **Supplementary Figure S13** shows that mQDs: DOPA offer more uptake than mQDs at days 0 and 7, and 14. This confirms that the conjugated mQDs: DOPA exhibits an increased uptake compared to mQDs alone; DOPA has a positive effect on the uptake of mQDs in differentiated neuron cells.

**Figure 7:**
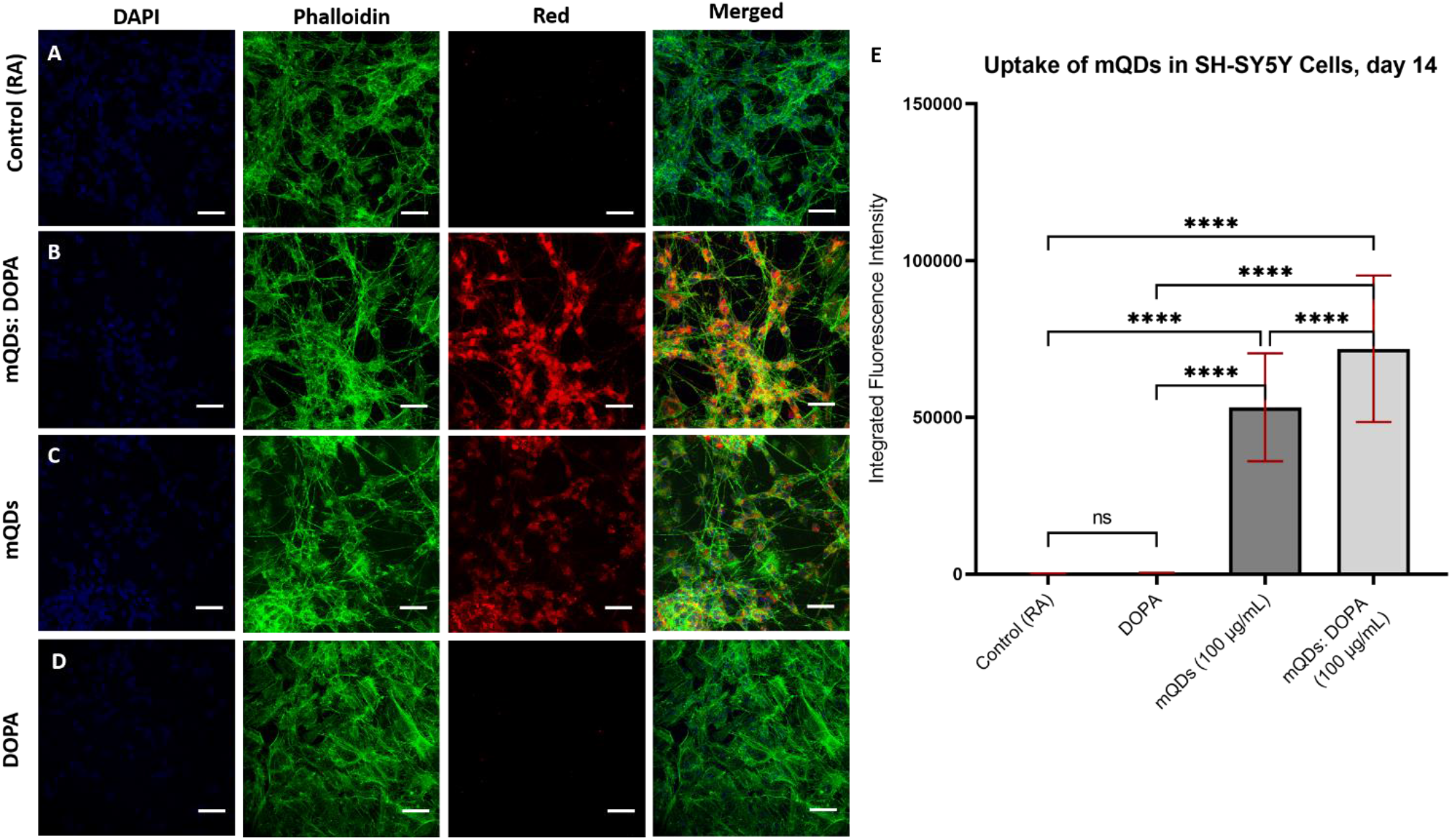
Dopamine enhances uptake of mQDs in differentiated SH-SY5Y neuroblastoma cells using RA. **(A)** Imaging of control SH-SY5Y cells (retinoic acid) with blue (DAPI labeled nucleus), green (Phalloidin labeled F-actin), and red (mQDs fluorescence) channels. **(B)** Imaging mQDs: DOPA–treated cells with blue, green, and red channels. **(C)** Imaging mQDs – treated cells with blue, green, and red channels. **(D)** Imaging of DOPA– treated cells with blue, green, and red channels. **(E)** Quantification of mQDs uptake in SH-SY5Y cells after 14 days.

To further test the potential of mQDs and mQDs-DOPA in collective cell migration, a scratch assay was performed on fully confluent slide of RPE-1 cells. The cells were then incubated and imaged with phase contrast microscopy at t=0 hours, t=12 hours, and t=24 hours to determine the percentage of wound closed with time (**Figure 8A, B, C**). After 12 and 24 hours in control, the wound gap remained around 79.5% and 72% of the original size, while DOPA showed no significant closure (**Figure 8D**). However, mQDs, for both 100 μg/mL and 200 μg/mL resulted in remarkable closure, lowering the gap size to 36.28% and 18.6% after 12 hours and 39.19% and 25.5% after 24 hours. In the case of mQDs: DOPA, after 12 hours and 24 hours of treatment, the 100 μg/mL closed gap to 53.03% to 40.7% of the original size, while the 200 μg/mL dose reduced size from 48.11% to 25.8% of original wound size (**Figure 8D**). The average length of the wounds was recorded and reported in **Supplementary Table S2 and** graphed in **Figure 8E**. Most notably, the mQDs: DOPA in 200 μg/mL decreased average wound gap from 407.6 μm to 105.3 μm in 24 hours, while the control only decreased wound size from 342.5 μm to 249.0 μm in 24 hours. Additionally, the mQDs in 200 μg/mL decreased wound size from 347.3 μm to 88.7 μm after 24 hours. We conclude that mQDs and mQDs: DOPA are both conducive to significant wound reduction (via increased cell migration) in RPE1 cells.

**Figure 8:**
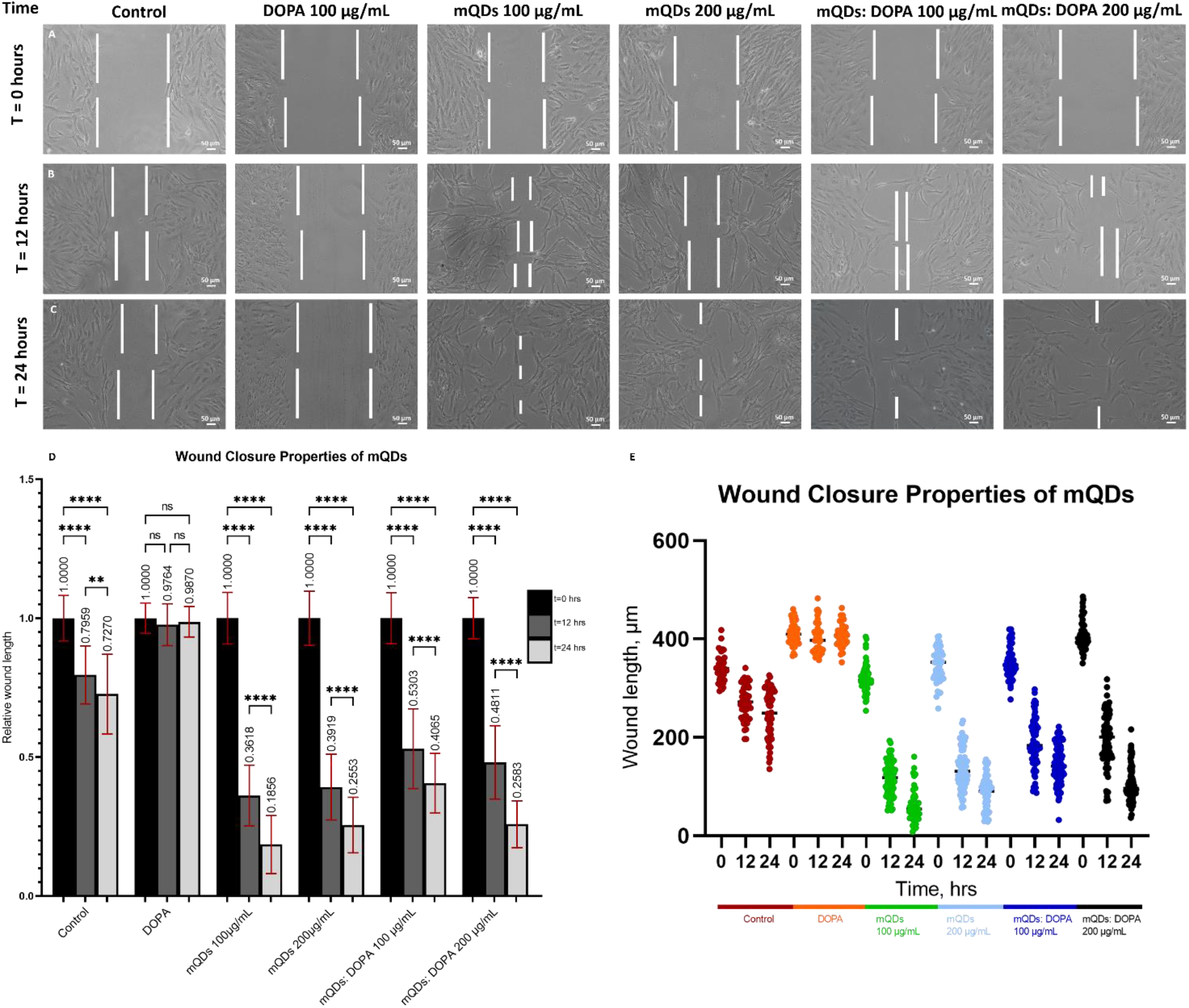
Wound healing in RPE-1 cells by mQDs and mQDs: DOPA. **(A)** RPE-1 cell imaging was taken at t=0 hours after the scratch was made with a pipette tip. (**B)** RPE-1 cell imaging was taken at t=12 hours. (**C)** RPE-1 cell imaging was taken at t=24 hours. Photos were taken using phase contrast microscopy imaging. (**D)** Graphed as a time function, relative wound length depends on the treatment applied to RPE-1 cells. (**E)** Absolute wound length, in μm, depends on the treatment applied to RPE-1 cells. The following thresholds were used when determining significance. ****: p< 0.0001. **: p< 0.01. ns: p>0.05 and denotes no significance.

## Conclusions and Discussions

The green synthesis using mango leaves reflux resulted in negatively charged red-emitting carbon quantum dots (mQDs), as shown in **Figure 1**. These mQDs were electrostatically conjugated with dopamine to form a bioconjugate (mQDs: DOPA), explained in **Figure 2**. The biocompatibility of mQDs and mQDs: DOPA was confirmed via MTT assay (**Supplementary Figure S5**). These results indicate that 200 μg/mL is an optimal dose for both mQDs and mQDs: DOPA. The endocytosis of mQDs was observed in live HeLa cells, where the mQDs invaded the cells in 2 minutes (**Figure 3**). The synthesized mQDs exhibited excellent red fluorescence and imaged the mouse tissues taken from the brain (**Figure 4A**), heart (**Supplementary Figure S6**), kidney (**Supplementary Figure S7**), and liver (**Supplementary Figure S8**). Various cells (SUM 159A and RPE-1, **Figures 3, 5**) and *in vivo* model system (zebrafish, **Figure 4E, S9**) were also successfully imaged with the mQDs and mQDs: DOPA. These results all indicated that the conjugation of mQDs with DOPA enhanced their cellular uptake in RPE-1 cells **Figure 5J**. The conjugation also resulted in mQDs: DOPA remaining inside SH-SY5Y cells for longer than mQDs alone (**Supplementary Figure S11**), which has implications for improved therapeutics and bio-imaging. As with many existing CQDs, the synthesized mQDs and mQDs: DOPA possess anti-cancer properties, passively reducing the viability of cancerous cells to around 40% of their original value (**Supplementary Figure S5C, S5D**). However, the improved uptake of mQDs: DOPA in SUM 159A makes them more efficient for tumor targeting and simultaneous bioimaging. Additionally, the cell death does not require a light trigger and subsequent phototherapy like current alternatives^18,19^, meaning it could be used in conjunction with other contemporary methods for an effective chemotherapy regime.

### MQDs

DOPA was also able to promote neuronal differentiation in SHSY-5Y cells, as demonstrated by increased dendrite length. Interestingly, the dendrite length is comparable to that of dendrites created using retinoic acid, a current standard for neuron differentiation (**Figure 6Q**). This behavior has applications for researching diseases caused by dysfunctional neurons or low dopamine, such as Parkinson’s disease^20^. Current treatments for these conditions focus on delivering dopamine agonists^21^ or transplanting human neural stem cells^22^, but studies have reported adverse side effects and high costs, respectively. mQDs: DOPA could act as an effective transport method for DOPA, triggering stem cell growth and differentiation and thus bypassing current treatment barriers. Another attractive characteristic of mQDs: DOPA, is their low cytotoxicity in RPE-1 cells. As our study showed, cell viability increased in the presence of mQDs and mQD:DOPA, indicating a potential proliferative nature. The proliferative effect was validated with the scratch assay in which mQDs (100 μg/m) and mQDs: DOPA (200 μg/m) showed wound size down to 18.53% and 25.83% of the original size, respectively, in **Figure 8D**. When treated with mQDs and mQDs: DOPA, wounds created in the scratch assay close much faster, exhibiting complete or near-full healing after 24 hours. An untreated scratch healed only around 30% in the same time frame (**Figure 8**). This indicates a positive effect of mQDs on RPE-1 cell migration. The ability to close wounds with both mQDs and mQDs: DOPA could be helpful in ointments due to cheap, sustainable, and easily scalable production processes.

The methodology reported in this paper has successfully created red-emitting mQDs derived from mango leaves capable of conjugation with dopamine. Naturally, the mQDs: DOPA has high fluorescent efficiency, and their red hue starkly stands out against commonly used blue/green dyes and imaging agents^23^. The DOPA conjugation also increases uptake within healthy cells, boosting imaging potential. Additionally, the mQDs: DOPA possess wound healing properties, as shown by their application within RPE-1 cells. mQDs: DOPA enhanced cell death within cancerous SUM159A cells while causing no reduction of benign cell viability. Most notably, however, the mQDs: DOPA were used to induce differentiation within SHSY-5Y neuronal cells, a novel finding. These combined results showcase mQDs: DOPA as versatile therapeutic/imaging agents capable of application to many contemporary healthcare issues including specific cellular and tissue targeting, uptake and subsequent use in bioimaging, diagnosis and therapy.

## Author contributions

PY and DB conceived the project and planned the experiments. Pankaj did the synthesis, characterization, cellular uptake, wound healing, microscopy of the brain tissue, and in vivo imaging and quantification. Dawson Benner conducted figure design/ statistical analysis, contributed data analysis of neuronal differentiation, and wrote the relevant sections of the manuscript. Landon Dahle designed the schematic figure and helped proofread the manuscript. Dr. Ritu Varshney conducted the neuronal differentiation experiment. Dr. Krupa Shah conducted the mouse tissue uptake experiments. Dr. Krupa Kansara conducted the uptake studies in zebrafish experiments and MTT assay. In addition to the authors, we thank the following for helping with specific experiments and infrastructure: Prof. Chinmay Ghoroi provided the lab facility for synthesizing mQDs. Akshant Kumawat, Vinod Morya, and Udisha Singh for helping with the spectroscopy and characterization techniques. Materials laboratory and Prof. Superb Mishra for DLS and FTIR. Prof. Saumykanti Khatua and Dr. Praneeth for decay time measurement.

## Acknowledgments

The authors sincerely thank all of the members of the D.B. group for critically reading the manuscript and for their valuable feedback. Gujarat University thanks Sync Bioresearch for providing mice to conduct experiments related to primary cells and tissues. P.Y. thanks IITGN-MHRD, GoI, Ph.D. fellowship; P.Y. acknowledges Director’s fellowship from IITGN for additional fellowship. K.K. thanks SERB, GoI, for National Postdoctoral Fellowship. K.S. acknowledges GSBTM and IITGN for the postdoctoral fellowship. D.B. thanks SERB, GoI for Ramanujan Fellowship, IITGN for the start-up grant, and Gujcost-DST, GSBTM, BRNS-BARC, and HEFA-GoI for research grants. D.B. is a member of the Indian National Young Academy of Sciences (INYAS). Imaging facilities of CIF at IIT Gandhinagar are acknowledged.

## References

1. Yang, H. et al. Dopaminergic Neuronal Differentiation from the Forebrain-Derived Human Neural Stem Cells Induced in Cultures by Using a Combination of BMP-7 and Pramipexole with Growth Factors. Front Neural Circuits 10, (2016).

2. Xiong, M. et al. Human Stem Cell-Derived Neurons Repair Circuits and Restore Neural Function. Cell Stem Cell 28, 112–126.e6 (2021).

3. Thompson, C. K. & Cline, H. T. Thyroid Hormone Acts Locally to Increase Neurogenesis, Neuronal Differentiation, and Dendritic Arbor Elaboration in the Tadpole Visual System. J Neurosci 36, 10356–10375 (2016).

4. Stevanovic, M. et al. SOX Transcription Factors as Important Regulators of Neuronal and Glial Differentiation During Nervous System Development and Adult Neurogenesis. Front Mol Neurosci 14, (2021).

5. Lam, H. J., Patel, S., Wang, A., Chu, J. & Li, S. In vitro regulation of neural differentiation and axon growth by growth factors and bioactive nanofibers. Tissue Eng Part A 16, 2641–8 (2010).

6. Björklund, A. & Dunnett, S. B. Dopamine neuron systems in the brain: an update. Trends Neurosci 30, 194–202 (2007).

7. John, V. L., Nair, Y. & Vinod, T. P. Doping and Surface Modification of Carbon Quantum Dots for Enhanced Functionalities and Related Applications. Particle & Particle Systems Characterization 38, 2100170 (2021).

8. Medintz, I. L. et al. Quantum-dot/dopamine bioconjugates function as redox coupled assemblies for in vitro and intracellular pH sensing. Nat Mater 9, 676–684 (2010).

9. Liu, Y. et al. Dopamine Receptor-Mediated Binding and Cellular Uptake of Polydopamine-Coated Nanoparticles. ACS Nano 15, 13871–13890 (2021).

10. Yang, Z. et al. Nitrogen-doped, carbon-rich, highly photoluminescent carbon dots from ammonium citrate. Nanoscale 6, 1890–1895 (2014).

11. Varshney, R., Gupta, S. & Roy, P. Cytoprotective effect of kaempferol against palmitic acid-induced pancreatic β-cell death through modulation of autophagy via AMPK/mTOR signaling pathway. Mol Cell Endocrinol 448, 1–20 (2017).

12. Zhu, Z. et al. red carbon dots: Optical property regulations and applications. Materials Today 30, 52–79 (2019).

13. Lan, M. et al. Two-photon-excited near-infrared emissive carbon dots as multifunctional agents for fluorescence imaging and photothermal therapy. Nano Res 10, 3113–3123 (2017).

14. Bao, X. et al. In vivo theranostics with near-infrared-emitting carbon dots—highly efficient photothermal therapy based on passive targeting after intravenous administration. Light Sci Appl 7, 91 (2018).

15. Maden, M. Retinoid signalling in the development of the central nervous system. Nat Rev Neurosci 3, 843–853 (2002).

16. Durston, A. J. et al. Retinoic acid causes an anteroposterior transformation in the developing central nervous system. Nature 340, 140–144 (1989).

17. Shipley, M. M., Mangold, C. A. & Szpara, M. L. Differentiation of the SH-SY5Y Human Neuroblastoma Cell Line. Journal of Visualized Experiments (2016) doi:10.3791/53193.

18. Hou, C., Chen, S. & Wang, M. Facile preparation of carbon-dot-supported nanoflowers for efficient photothermal therapy of cancer cells. Dalton Transactions 47, 1777–1781 (2018).

19. Zhang, Y. et al. Carbon dots nanophotosensitizers with tunable reactive oxygen species generation for mitochondrion-targeted type I/II photodynamic therapy. Biomaterials 293, 121953 (2023).

20. Cramb, K. M. L., Beccano-Kelly, D., Cragg, S. J. & Wade-Martins, R. Impaired dopamine release in Parkinson’s disease. Brain (2023) doi:10.1093/brain/awad064.

21. Fahn, S. The history of dopamine and levodopa in the treatment of Parkinson’s disease. Movement Disorders 23, S497–S508 (2008).

22. Dhara, S. K. & Stice, S. L. Neural differentiation of human embryonic stem cells. J Cell Biochem 105, 633–640 (2008).

23. Yadav, P. et al. Tissue-Derived Primary Cell Type Dictates the Endocytic Uptake Route of Carbon Quantum Dots and In Vivo Uptake. ACS Appl Bio Mater 6, 1629–1638 (2023).

