## Supplementary material for "Dopamine functionalized, red carbon quantum dots for *in vivo* bioimaging, cancer therapeutics, and neuronal differentiation": Supplimentary file

### 1. Materials and Methods :

#### 1.1 Materials:

Mango leaves were obtained from mango trees at IIT Gandhinagar campus. Silicon oil was purchased from X-chemicals, ethanol (>99.9%) from Changshu Hongsheng Fine Chemicals Co. Ltd. The filter of 0.22 micrometer was purchased from Merck, as well as the deionized water was taken from Merck Millipore. Phalloidin and dopamine hydrochloride were purchased from Sigma Aldrich. The cell culture dishes, dimethyl sulfoxide (DMSO) and rhodamine B were purchased from Himedia. DMEM (Dulbecco's modified Eagle's medium), DMEM F12 (1:1), FBS (fetal bovine serum), and trypsin-EDTA (0.25%) were obtained from Gibco. Retinoic acid was purchased from Sigma-Aldrich. All the purchased chemicals were of analytical grade without the need for further purification.

#### 1.2 Synthesis of mQDs:

Mango leaves were plucked from a mango tree on IIT Gandhinagar's campus. The leaves were washed and dried under shade naturally. Then, leaves were crushed using a kitchen grinder to make powder. The powder was then dissolved in ethanol in 1:10 (w/v) ratio using a magnetic stirrer under a constant stirring (300 rpm) for 4 hours . The dissolved powder is then centrifuged at 10,000 rpm for 10 minutes at room temperature to get the mango leaf extract. The extract is then refluxed for 2 hours at 160$^{o}$C and then cooled down naturally to obtain mQDs. The formed mQDs were then filtered using a 0.22 µm syringe filter. Finally, the solvent was evaporated using rotavapor to obtain powdered mQDs. The mQDs powder is then characterized and further used to study their cellular uptake, tissue bioimaging.

#### 1.3 Characterization of mQDs:

The optical properties of mQDs, both UV-Vis absorbance and fluorescence emission were recorded using Spectrocord-210 Plus Analytokjena (Germany) and FP-8300 Jasco spectrophotometer (Japan) respectively.

The sample preparation for AFM was done on a freshly peeled mica sheet. A 5µL drop of powdered mQDs (1mg/mL in water ) and mQDs: DOPA (1mg/mL in water ) was dropped on a mica sheet. The mica sheet was then kept in a desiccator for drying. Finally, the AFM imaging was done in tapping mode using the Bruker AFM instrument.

FTIR spectra of powdered mQDs, mQDs: DOPA, DOPA and mango leaves were recorded from 400 cm$^{-1}$ to 4000 cm$^{-1}$ using spectrum 2 PerkinElmer in ATR mode. The powder XRD diffraction pattern was recorded using Bruker-D8 Discover with a speed of 0.2/min from 5° to 90°.

Dynamic Light Scattering (DLS) was done by using the Malvern analytical Zetasizer Nano ZS for the solution-based size characterization of mQDs and mQDs: DOPA. The data was then plotted in OriginPro software using Gaussian fit.

For lifetime analysis, the mQDs powder was dissolved in deionized water. Then, the sample(10 mg/mL) was prepared by drop casting on a coverslip and allowed to dry. The data was recorded using an LDH-D-C-488 nm pulsed laser source (Pico Quant GmbH), detected by SPAD (Excelitas) connected to TimeHarp 260 board (Pico Quant GmbH).

The photostability of mQDs under constant UV treatment was measured for 50 minutes. mQDs dissolved in ethanol (1 mg/mL) were kept in an UV transilluminator, and the fluorescence intensity at 670 nm was recorded every 10 minutes upon excitation with 400 nm. The relative fluorescence intensity was plotted by dividing each fluorescence intensity by the maximum fluorescence intensity.

The mQDs and mQDs: DOPA were dissolved in serum-free media (SFM) after lyophilization for bio imaging in cells, mouse tissues, zebrafish and neuronal cells.

**1.4 Quantum Yield Calculation:**

The quantum yield (Φ) of the mQDs was calculated using rhodamine B as reference. For calculation of quantum yield, a solution of concentration of 1 mg/mL of rhodamine B (RB) was made in milli-Q water. 0.1 mL of this 1 mg/mL was taken and dissolved into 1 mL of milli Q so that final concentration of RB was 100 µg/mL. The absorbance and fluorescence were taken by taking 2 µl of RB in the cuvette, keeping the final volume to 3 mL . The next reading was found by taking 4 µL of RB, keeping the final volume to be 3 mL. Similarly, the subsequent readings were found by increasing the concentration of RB by 2 µL and keeping the final volume to 3 mL.

The mQDs were dissolved in ethanol (99.5%) having concentration of 1mg/mL. The absorbance and fluorescence were found by taking 2 µL of mQDs in the cuvette keeping the final volume to 3 mL. The subsequent readings were taken by increasing the concentration of mQDs by 2 µL keeping the final volume to 3 mL.

The absorbance for both RB and mQDs were kept less than 0.1 at 527 nm and 670 nm respectively. RB (literature Φ = 0.31) was dissolved in water (refractive index = 1.33) and mQDs were dissolved in ethanol (refractive index = 1.36) . Their fluorescence spectra were recorded at the same excitation of 400 nm. Then, by comparing the integrated photoluminescence intensities (excited at 400 nm) and the absorbance values of mQDs with the reference RB, quantum yield of the mQDs was determined. The data was plotted (**Figure S3**) and the slopes of the sample (mQDs) and the standards (RB) were determined. The data showed good linearity having mean square deviation 0.99 (reference RB) and 0.99 (sample mQDs).

The quantum yield was calculated using the below equation

$$\Phi_{mQDs} = \Phi_{RB} \times ( m_{QDs} / m_{RB}) \times (\eta^2_{mQDs} / \eta^2_{RB}),$$

where Φ is the quantum yield, m is slope, η is the refractive index of the solvent, RB is the reference and mQDs is the sample. The quantum yield for mQDs was found to be 202% relative to RB.

### 1.5 Loading of DOPA on mQDs:

The loading of DOPA on mQDs was calculated by plotting a standard curve of absorbance of DOPA with increasing concentration **as shown in figure S4B.** The linear equation of absorbance with concentration was $\boldsymbol{y} = 0.04829\boldsymbol{x} + 0.00825$, where $\boldsymbol{y}$ represents absorbance and $\boldsymbol{x}$ represents concentration of dopamine in mg/mL.

The mQDs and DOPA were taken in ratio of 1:2 (0.5mg : 1mg ) and then were mixed in water as solvent and kept overnight at 4°C. After 12 hours mQDs and mQDs: DOPA(1:2) were dialyzed using 10kDA snakeskin membrane for 24 hours. The absorbance of the solvent (water) was recorded for both mQDs and mQD:DOPA. The absorbance values for mQDs and mQD:DOPA were 0.0101 and 0.0413 respectively as shown in **figure S4C.** Since the mQDs might have leaked from the dialysis bag, their absorbance was subtracted to calculate the actual loading of DOPA. The loading of DOPA on mQDs was calculated to be 52.47%.

$\boldsymbol{y} = 0.04829\boldsymbol{x} + 0.00825$

$(0.0413 - 0.0101) = 0.04829\boldsymbol{x} + 0.00825$

$0.0312 = 0.04829\boldsymbol{x} + 0.00825$

$0.0312 - 0.00825 = 0.04829\boldsymbol{x}$

$0.02295 \div 0.04829 = \boldsymbol{x}$

0.47525 mg/mL = concentration of DOPA in solvent .

0.52475 mg/mL = concentration of DOPA in dialysis bag.

### 1.6 Biocompatibility Assay:

In order to assess the cytotoxicity of mQDs, and mQDs: DOPA approximately 1×105 /100µl of SUM 159, RPE1 (Retinal Pigment Epithelial Cell Line) and SH-SY5Y cells were seeded in 96-well in media (Ham's F-12 complete media for SUM159 ,DMEM containing 10% FBS, 1x antibiotic (Pen strep) for RPE1 ) and DMEM F12 (1:1) complete media respectively. The cells were allowed to acclimatized for 24 hours in a 5% CO2 at 37ºC before the experiment. Next day, cells were washed with PBS and treated with increasing concentrations of mQDs and mQDs: DOPA (100, 200, 300, 400, 500 µg/ml) for 24 hours in a serum free medium. After the treatment ,culture medium was discarded, 100 µl of media (Ham's F-12 serum free media for SUM159 and DMEM complete media for RPE1) containing MTT (5mg/ml) was added, incubated for 3-4 hours to determine the mitochondrial dehydrogenase activity of viable cells. Media was aspirated again, and 100 µl of DMSO was added to each well to dissolve the purple formazan crystals and absorbance spectra was measured at 570 nm using Multiskan microplate reader. Experiment was done in triplicate, normalized to corresponding well containing DMSO whereas, non-treated mQDs well were considered as control to calculate the % cell viability of each well.

Cell Viability (%) = OD (Sample) – OD (Blank)/ OD (Control) – OD (Blank) ×100

### 1.7 Cellular Uptake

The cellular uptake of mQDs was studied in SUM 159 and RPE1 cells. The SUM 159 cells were cultured and maintained in Ham's F12 complete media in T 25 flask. Similarly, RPE1 cells were cultured and maintained in DMEM complete media in T 25 flask. The experiment was performed by seeding 105 cells per well on 10mm glass coverslip in a 24 well plate, 24 hours before the experiment. Prior to the experiment cells were checked under a microscope to visualize their attachment and spreading. The cells were washed 3 times with 1X PBS and then incubated with Hams serum free media for 15 minutes. The cells were treated with increasing concentrations of mQDs (0 ug/ml, 50 ug/ml 100 ug/m, 200 ug/ml, 300 ug/ml). After the treatment, the cells were incubated at 37$^{o}$C for 20 minutes. The excess surface bound mQDs were removed by washing 3 times with 1X PBS. The cells were fixed using 4% PFA ( paraformaldehyde) for 15 minutes at 37$^{o}$C. The fixed cells were again washed three times with 1X PBS . Finally, the coverslips were mounted on a glass slide using mowiol containing Hoechst to mark the nucleus.

**1.8 Uptake assay of mQDs using in differentiated neurons using RA :**

Neuronal differentiation was performed as per previous protocol. The experiment was performed by seeding $10^5$ cells per well on 10mm glass coverslip in a 4 well plate, 24 hours before the experiment.

1. Differentiation media 1 (DFM1) was added on day 0,3,5,7.
2. Differentiation media 2 (DFM2) was added on day 8, 10.
3. Differentiation media 3 (DFM3) was added on day 11, 14.

The cells were fixed on day 0, 7 and 14 of RA treatment for bioimaging. The protocol was followed as mentioned below:

The experiment cells were checked under a microscope to visualize their attachment and spreading. The cells were treated with mQDs (100 µg/mL), mQDs: DOPA (100 µg/mL) and DOPA (50 µg/m). After the treatment, the cells were incubated at 37$^{o}$C. The same treatment was repeated on day 3 and day 9. The cells were under continuous observation for 10 days. On day 10 the excess surface bound mQDs were removed by washing 3 times with 1X PBS. The cells were fixed using 4% PFA ( paraformaldehyde) for 15 minutes at 37$^{o}$C. The fixed cells were again washed three times with 1X PBS .Phalloidin treatment was given for 15 minutes. The fixed cells were again washed three times with 1X PBS. Finally, the coverslips were mounted on a glass slide using mowiol containing Hoechst to mark the nucleus.

| Media Type | Component | Volume |
|---|---|---|
| **DFM1(10mL)** | | |
| | DMEM F12(1:1) | Make upto final volume |
| | FBS | 250ul |
| | Pen-strap | 100ul |
| | RA(5mM stock) | 20ul |
| **DFM2(10mL)** | | |
| | DMEM F12(1:1) | Make upto final volume |
| | FBS | 100ul |
| | Pen-strap | 100ul |

| | | |
|---|---|---|
| | RA(5mM stock) | 20ul |
| **DFM3 ( 10mL)** | Neurobasal Media | Make up to final volume |
| | KCl(1M stock) | 200ul |
| | Pen-strap | 100ul |
| | Glutamaxl(100X stock) | 100ul |
| | B 27 supplement | 200ul |
| | RA (5mM stock) | 20ul |

**Table S1: Components of different media used in the differentiation of SH-SY5Y**

**1.9 Neuronal differentiation of SH-SY5Y by mQDs: DOPA:**

The experiment was performed by seeding $10^5$ cells per well on 10mm glass coverslip in a 4 well plate, 24 hours before the experiment. On day 0 of the experiment cells were checked under a microscope to visualize their attachment and spreading. The cells were treated with mQDs (100 µg/mL), mQDs: DOPA (100 µg/mL) and DOPA (50 µg/mL). After the treatment, the cells were incubated at 37$^{o}$C. The same treatment was repeated on day 3 and day 9. The cells were under continuous observation for 10 days. On day 10 the excess surface bound mQDs were removed by washing 3 times with 1X PBS. The cells were fixed using 4% PFA ( paraformaldehyde) for 15 minutes at 37$^{o}$C. The fixed cells were again washed three times with 1X PBS .Phalloidin treatment was given for 15 minutes. The fixed cells were again washed three times with 1X PBS. Finally, the coverslips were mounted on a glass slide using mowiol containing Hoechst to mark the nucleus.

1. The cells were seeded for 24 hours before experiment in 4 well plate
2. Then treatment of mQDs, mQDs: DOPA , DOPA , RA (10uM) was given on day 0, day 3, day 9.
3. In case of mQDs: DOPA the cells were healthy until day 2; however, cells were observed to be stressed on day 4. The treatment media was removed, and then fresh serum free media was added. The cells were healthy again on day 5.
4. The complete media (DMEM:F12(1:1)) was changed on every even day.
5. Continuous monitoring was done by taking bright field images on each day.

**1.10 *Ex vivo* Cellular Uptake of mQDs in mouse kidney, heart and liver tissues sections:**

The protocol to isolate kidney, heart and liver sections from mice are described above. From the isolated kidney, heart and liver tissues, small sections were extracted and washed with cold PBS, immediately, tissues were snap frozen using liquid nitrogen and cut into 1 mm of small slices using sterile scalpel. Once cut, tissue slices were allowed to revive in prepared medium DMEM:F12 (1:1) for 20 min in a 5% $CO_2$ at 37ºC from the shock of snap freezing. Next, the tissues were incubated in PBS for 5 min and washed twice, cut into further small slices using scalpel and incubated with mQDs (50, 100, 200 mg/mL) for 60 min in a 5% $CO_2$ at 37ºC in a serum free media (DMEM+F12). After incubation, slices were washed thrice with PBS to remove extra mQDs and fixed with fixative solution (4% PFA) for 15 min at 37ºC. Once tissue slices were fixed, washed with PBS and mounted with Mowiol containing Hoechst for further confocal imaging.

**1.11 Zebrafish Maintenance and Ethical Approval:**

1-day old Swiss albino (*Mus musculus*) neonatal mice were used in study. Following conditions were kept to maintain the zebrafish. Mice pups were kept in a clean environment with 12 hours light/12 hours dark cycle conditions. The air was conditioned at 21±3 ºC and the relative humidity was maintained between 30-70% with 100% exhaust facility. Institutional ethical approval was obtained for all the zebrafish experiments conducted in the study (SBR/M3/008/2022).

**1.12 Zebrafish husbandry and maintenance**

The Assam wild type zebrafish were acquired from local vendors and were maintained at controlled laboratory conditions in Ahmedabad University according to the method of Kansara et al., 2019. The male and female fishes were retained in 20 L tanks with internal conditions maintained as per ZFIN in artificially prepared fresh water. The quality of the water in the aquarium were checked regularly for the pH (6.8-7.4), conductivity (250-350 mg/L), TDS (220-320 mg/L), salinity (210-310 mg/L) and dissolved oxygen (>6 mg/L), using multi-parameter instrument (Model PCD 650, Eutech, India). The strips purchased from Macherey-Nagel Inc. (USA) were used to check the contents of ammonia (QUANTOFIX®; Reference number 90714) and nitrate/nitrite (QUANTOFIX®; Reference number 91313). The photoperiod was maintained to 14 h light/10 h dark cycle at 26-28$^0$C. The zebrafish were supplemented with brine shrimp daily. Plastic traps for embryo collection were set up in the evening with the ratio of 3 females and 2 males. For experiments, embryos were collected and incubated in E3 medium (5 mmol/L NaCl, 0.17 mmol/L KCl, 0.33 mmol/L $CaCl_2$, and 0.33 mmol/L $MgSO_4$, dissolve in $dH_2O$, adjust pH at 7.2 and autoclave E3 media; store it at room temperature) at 28˚C.

**1.13 *In vivo* uptake of mQDs and mQDs: DOPA in Zebrafish Model:**

*In vivo* uptake assays were performed according to organization for economic cooperation and development (OECD) guidelines. At 72 hpf (hours post fertilized), dead larva was removed and the remaining larva placed in six-well plates (Corning, NY, USA) with 15 larvae in each well. Two groups of larvae were treated with mQDs and mQDs: DOPA at concentrations of 200 µg/mL each and incubated for 4 hours. One well designated as control in each group without nanoparticles. Post treatment, the medium was replaced with fresh E3 media and larva were washed to remove the excess mQDs and fixed with fixative solution (4% PFA) for 2 minutes. Post fixation, the larva was mounted with mounting solution Mowiol and allowed to dry for further confocal imaging analysis.

#### 1.14 Confocal Imaging and Processing:

The confocal imaging of fixed cells (63x oil immersion) and fixed tissues/embryos (10x) was performed using Leica TCS SP8 confocal laser scanning microscope (CLSM, Leica Microsystem, Germany). Different fluorophores were excited with different lasers i.e., for Hoechst (405 nm),phalloidin (488 nm), mQDs (633 nm). The pinhole was kept 1 airy unit during imaging. Image quantification analysis was performed using Fiji ImageJ software. For the quantification analysis, whole cell intensity was measured at maximum intensity projection, background was subtracted and measured fluorescence intensity was normalized against unlabeled cells. A total of 40-50 cells were quantified from collected z-stacks for each experimental condition.

#### 1.15 Statistical Analysis:

Statistical analysis was performed using Graph Pad Prism software (version 8.0.2). All the data were expressed as means ± standard deviation (SD) or means ± standard error from two independent experiments. p values were calculated using one-way ANOVA and two -tailed unpaired student t-tests with 95% confidence interval.

#### 1.16 Scratch Assay Procedure

Six groups of cells were plated in DMEM serum-free media: control, DOPA 100 µg/mL, mQDs 100 µg/mL, mQDs 200 µg/mL, mQDs: DOPA 100 µg/mL, and mQDs: DOPA 200 µg/mL. The control was plated with just the cells, and the DOPA with just dopamine, in order to test whether the mQDs, the DOPA, or the mQDs: DOPA contributed the wound closure effect. Each cell plate was scraped straight up and down with a pipette tip and gently washed with PBS to remove detached cells. Afterwards, the cells were incubated and imaged with phase contrast microscopy at t=0 hours, t=12 hours, and t=24 hours to determine how wounds progressed as time passed. Measurements were made using ImageJ and compared to original scratch width.

**Supplementary Figures:**

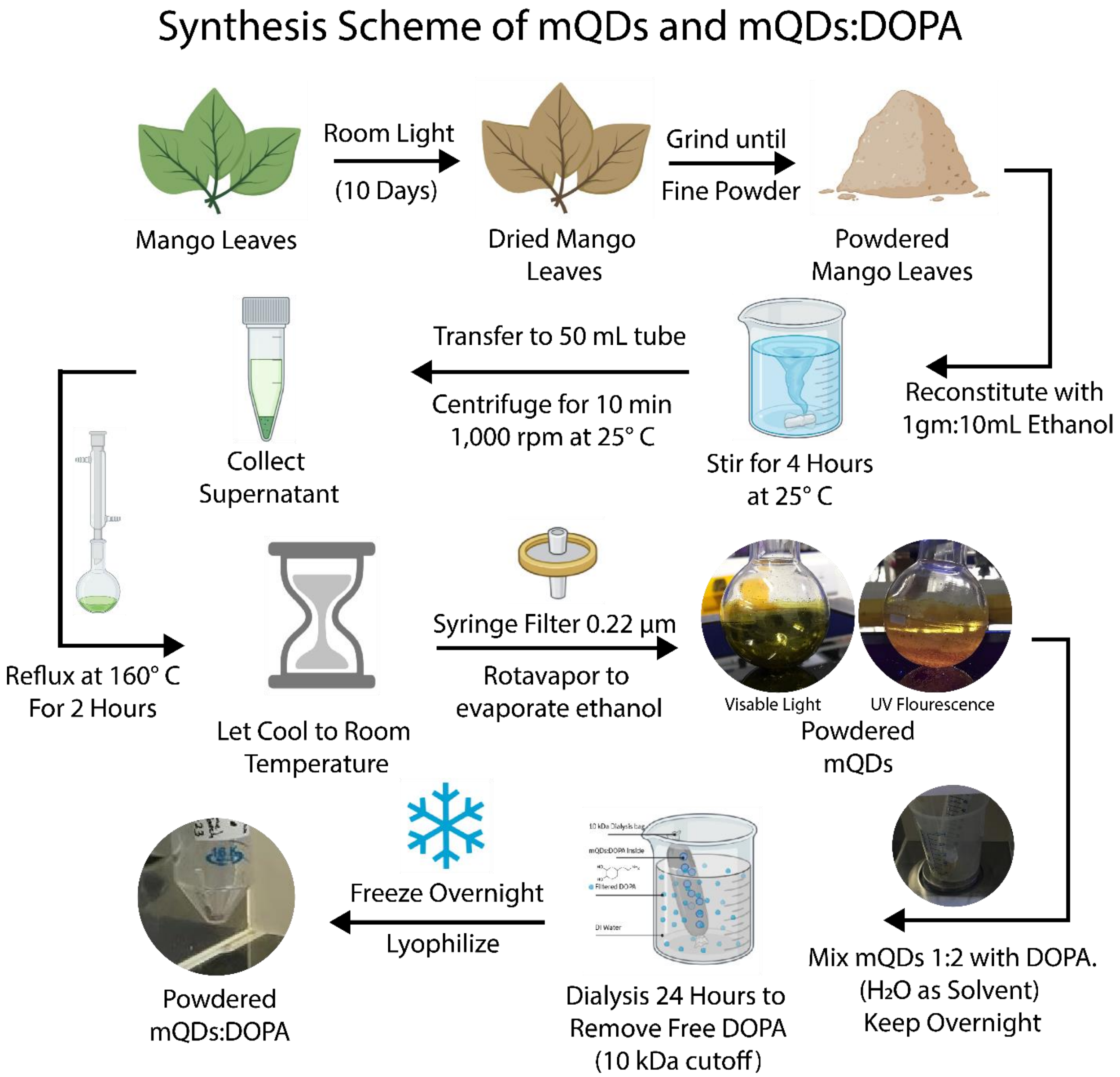

**Scheme 1:** Synthesis of mQDs and conjugation with DOPA.

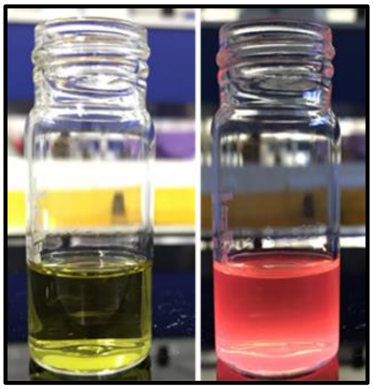

**Figure S1:** mQDs solution under room light (left) and UV light (right).

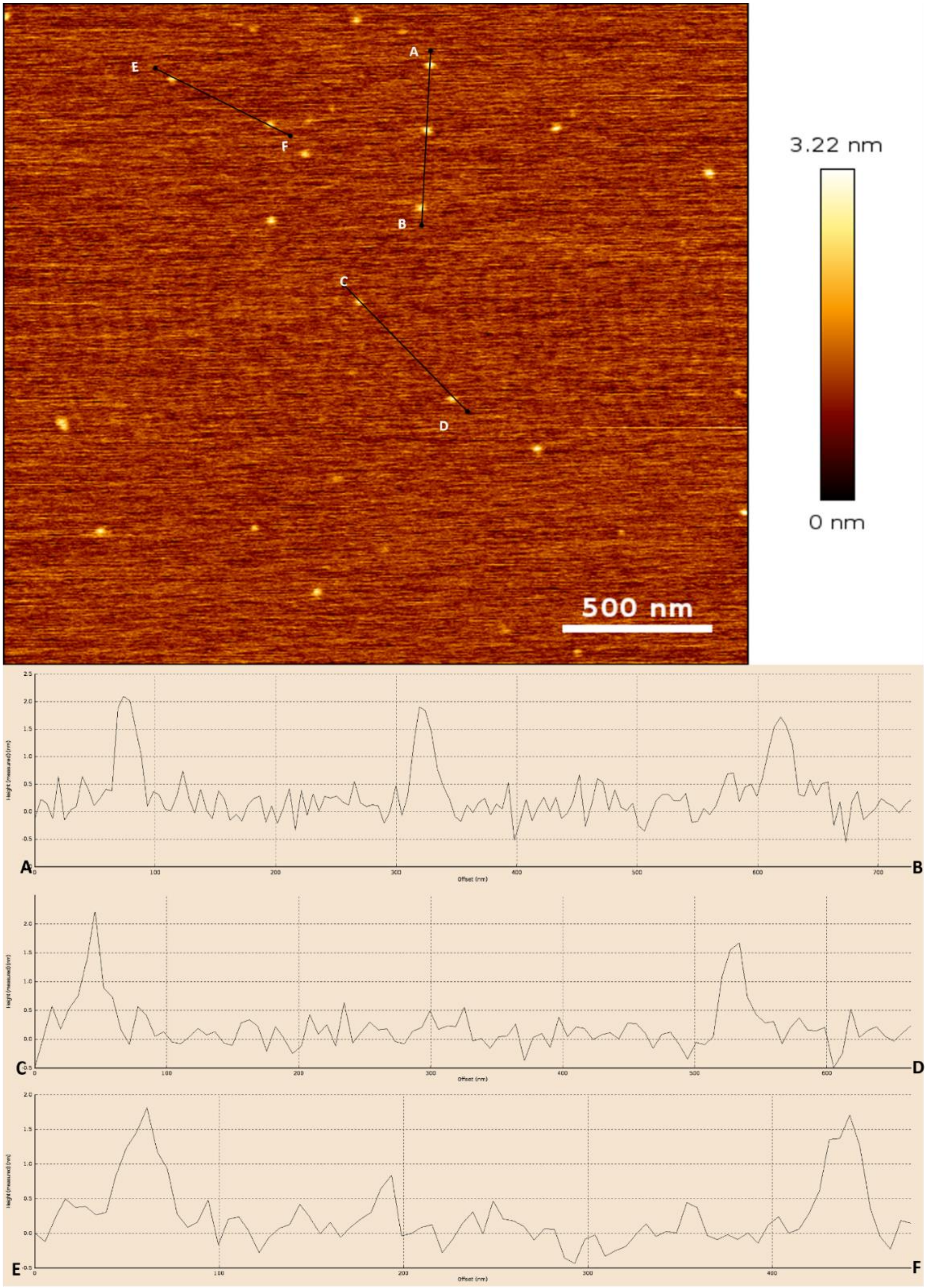

**Figure S2: Height profile of mQDs.** The lines AB, CD and EF gives the topographic height profile of the surface. The height of mQDs is 2 nm.

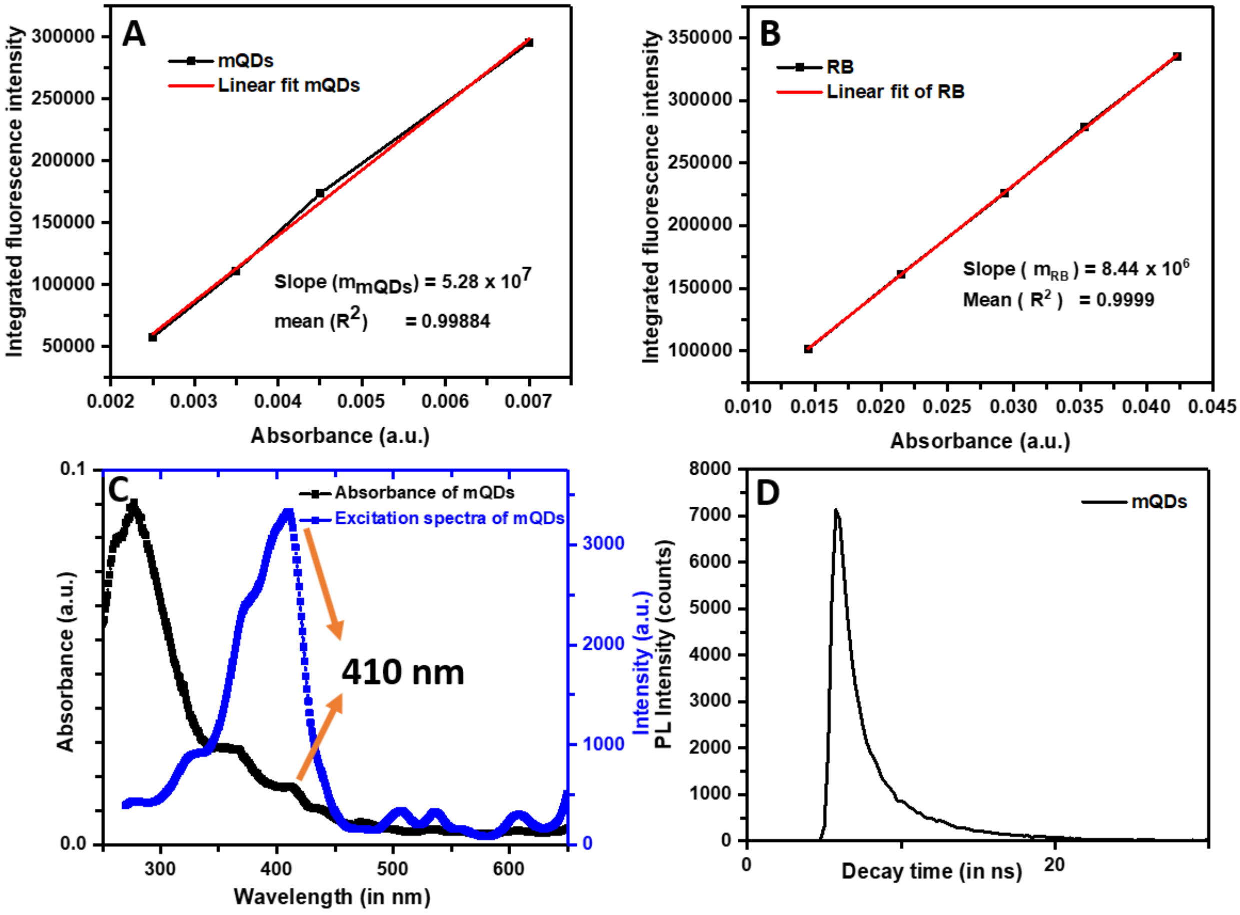

**Figure S3: Quantum yield quantification and calculation. (A)** Integrated fluorescence intensity vs absorbance for mQDs. **(B)** Integrated fluorescence intensity vs absorbance for Rhodamine B standard. **(C)** UV-visible absorbance spectra of mQDs. the peak at 410 nm overlaps with the excitation maxima **(D)** Excitation spectrum of mQDs upon emission with 670 nm, the excitation maxima is 410 nm.

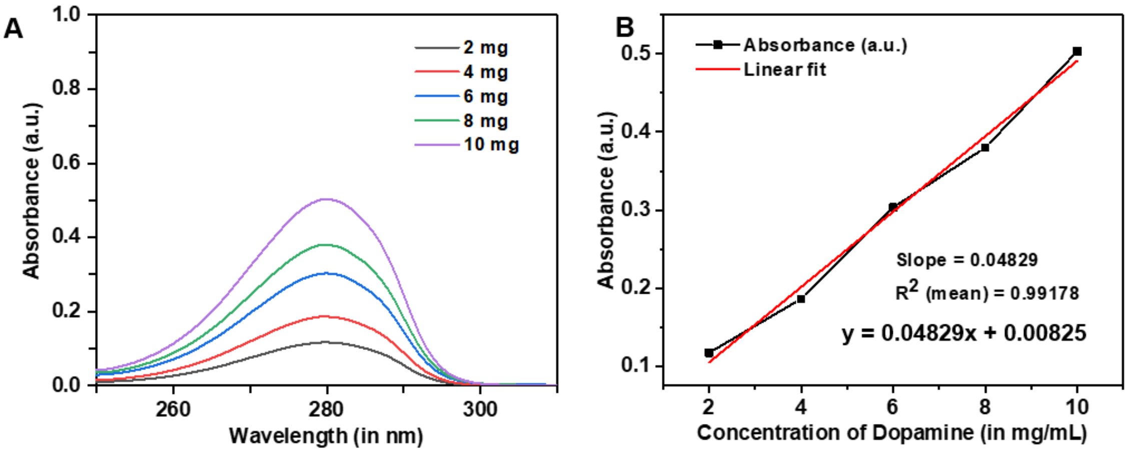

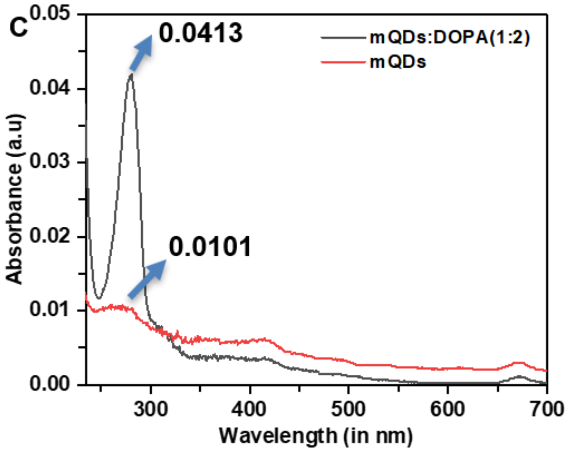

D

| Components | Zeta Potential (mV) |
|---|---|
| mQDs (0.5mg/mL) | -32.4 |
| mQDs:DOPA (1:2) | -21.5 |
| mQDs:DOPA (1:10) | -6.39 |
| mQDs:DOPA (1:20) | +14.5 |
| DOPA (10 mg/mL) | +7.96 |

**Figure S4: Loading of DOPA on mQDs. (A)** Absorbance at 280 nm with increasing concentration of DOPA. **(B)** Linear standard plot of absorbance at 280nm verses concentration of DOPA. **(C)** Absorbance of solvent of mQDs and mQDs: DOPA at 280 nm after 24 hours of dialysis. **(D)** Zeta potential of mQDs, DOPA and mQDs: DOPA (1:2, 1:10, and 1:20)

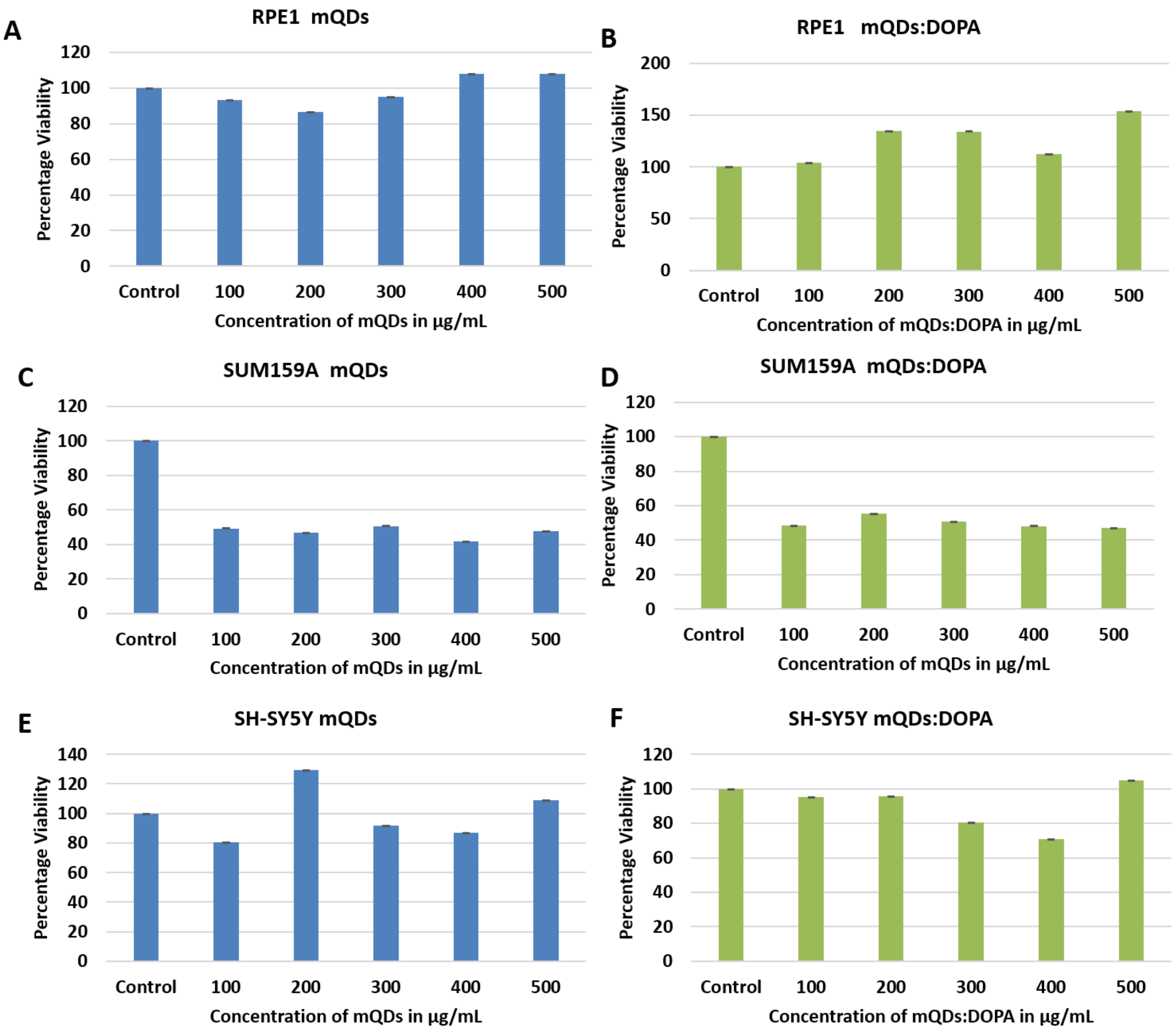

**Figure S5: RPE-1, SUM159A, and SH-SY5Y Cell Viability with varying mQDs and mQDs: DOPA dosages. (A)** RPE-1 cell viability with varied mQDs dosages from 0 to 500 µg/mL. **(B)** RPE-1 cell viability with varied mQDs: DOPA dosages from 0 to 500 µg/mL. **(C)** SUM159A cell viability with varied mQDs dosages from 0 to 500 µg/mL. **(D)** SUM159A cell viability with varied mQDs: DOPA dosages from 0 to 500 µg/mL. **(E)** SH-SY5Y cell viability with varied mQDs dosages from 0 to 500 µg/mL. **(F)** SH-SY5Y cell viability with varied mQDs: DOPA dosages from 0 to 500 µg/mL.

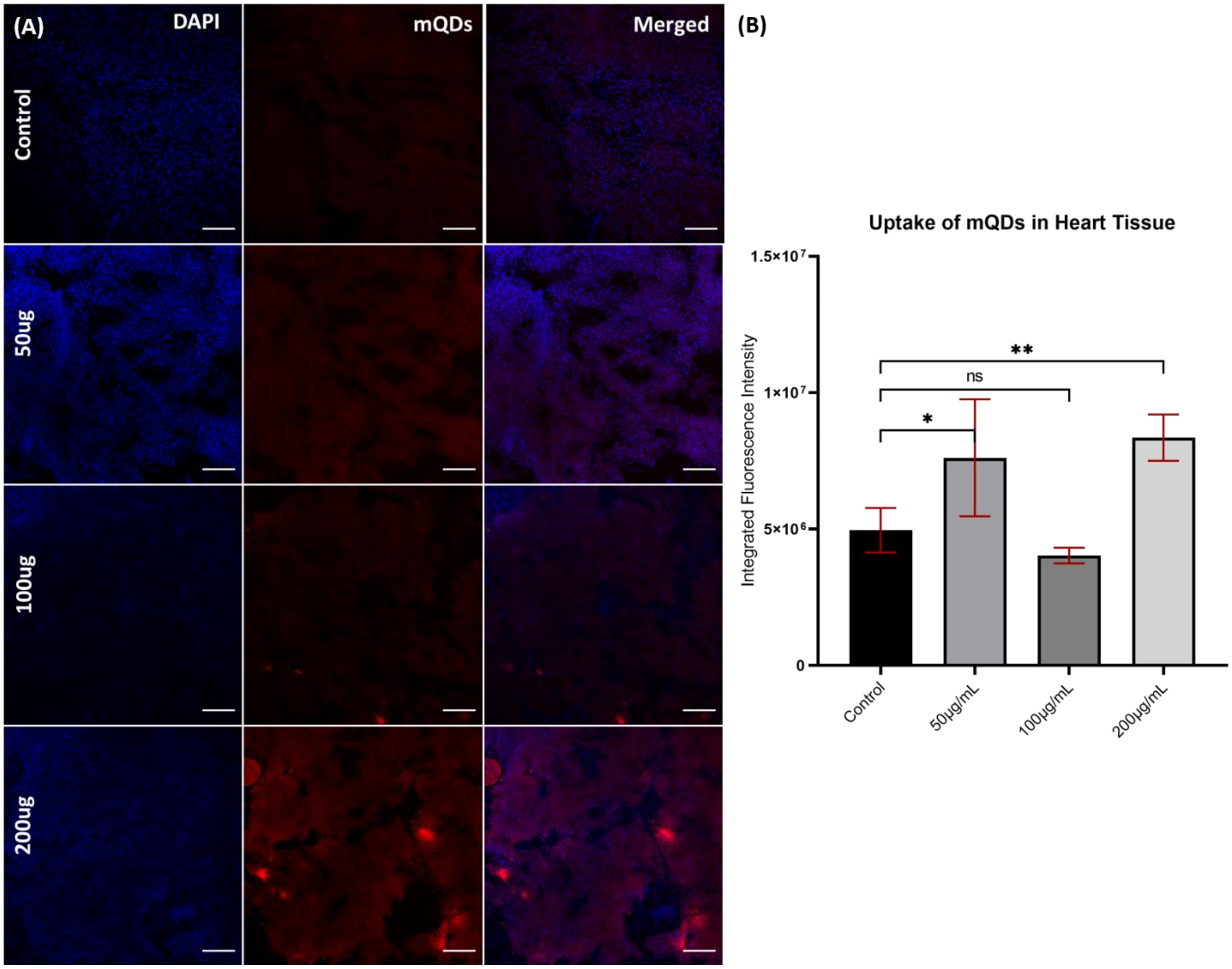

**Figure S6: mQDs uptake in heart tissue. (A)** Imaging results of control, 50 µg/mL, 100 µg/mL, and 200 µg/mL, with blue (DAPI labelling nucleus) and red (mQDs fluorescence) channels. **(B)** Quantified uptake of mQDs in heart tissue at differing doses. Scale bar 5 µm. The following thresholds were used when determining significance. **: P< 0.01. *: P< 0.05. ns: P>0.05 and denotes no significance. (n = 5 tissues sections for quantification)

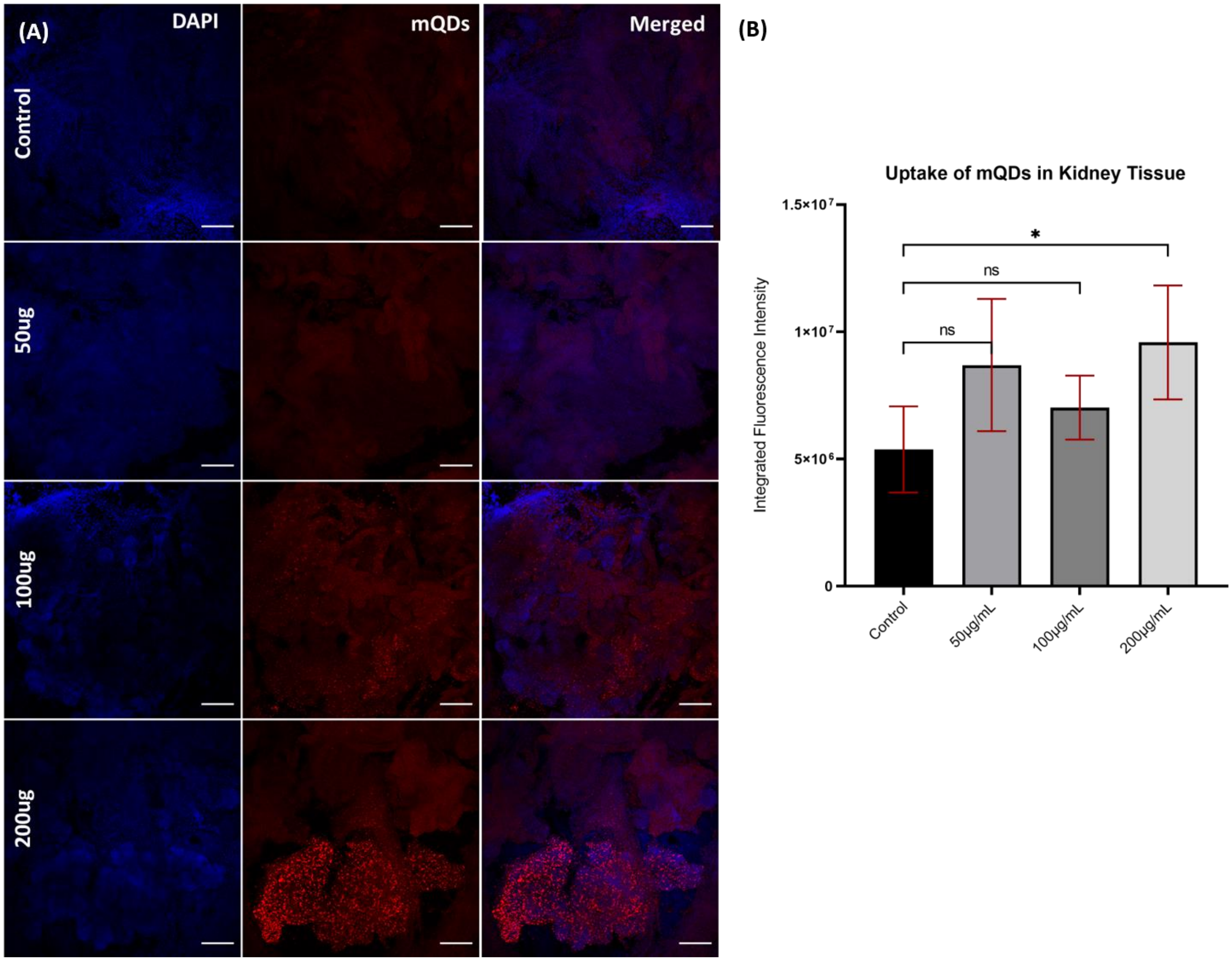

**Figure S7: mQDs uptake in kidney tissue. (A)** Imaging results of control, 50 µg/mL, 100 µg/mL, and 200 µg/mL with blue (DAPI labelling nucleus) and red (mQDs fluorescence) channels. **(B)** Quantified uptake of mQDs in kidney tissue at differing doses. Scale bar 5 µm. The following thresholds were used when determining significance. *: P< 0.05. ns: P>0.05 and denotes no significance. (n = 5 tissues sections for quantification)

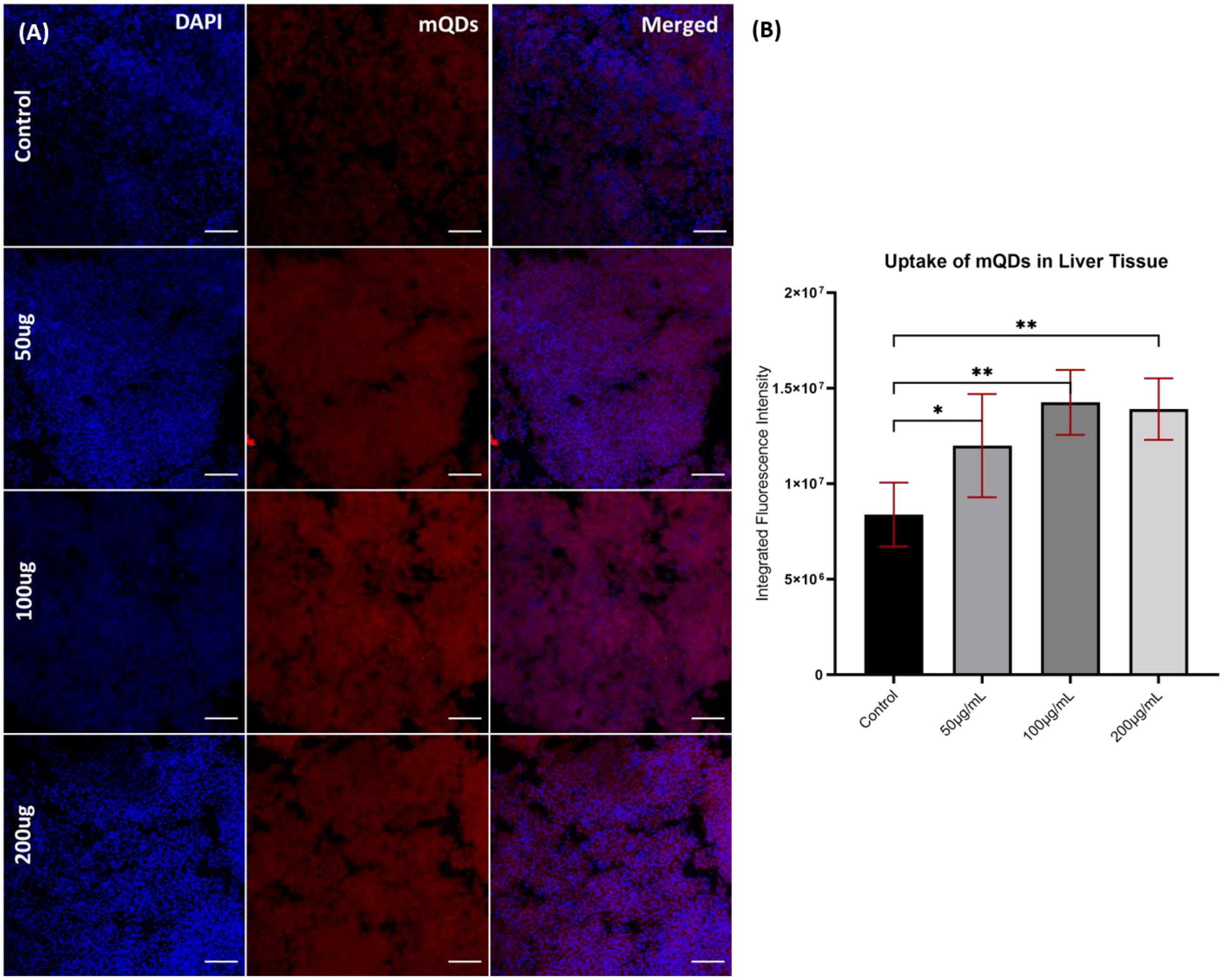

**Figure S8: mQDs uptake in liver tissue. (A)** Imaging results of control, 50 µg/mL, 100 µg/mL, and 200 µg/mL with blue (DAPI labelling nucleus) and red (mQDs fluorescence) channels. **(B)** Quantified uptake of mQDs in liver tissue at differing doses. Scale bar 5 µm. The following thresholds were used when determining significance. **: P< 0.01. *: P< 0.05. (n = 5 tissues sections for quantification)

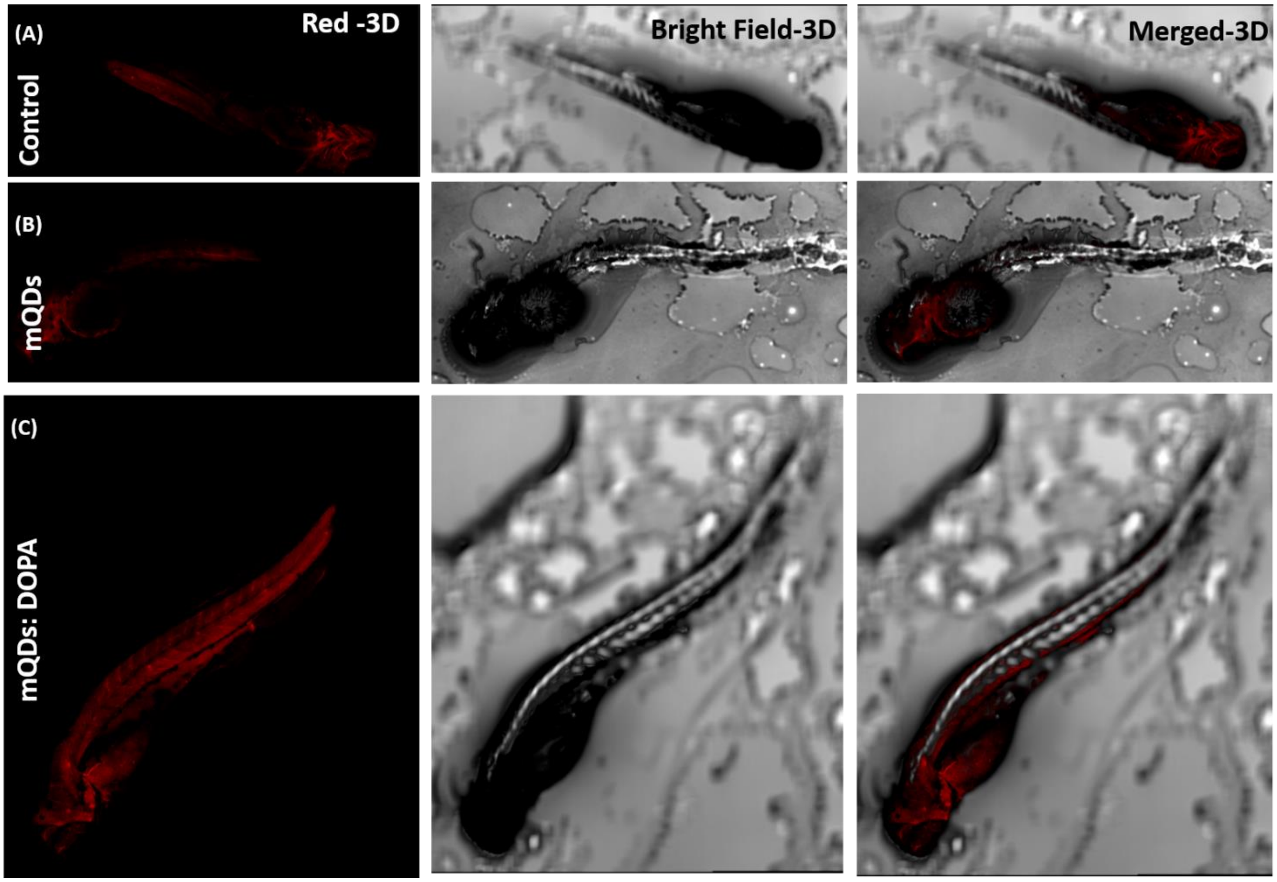

**Figure S9: uptake of mQDs and mQDs: DOPA in zebrafish (A)** The zebrafish without treatment **(B)** Zebrafish treated with mQDs (200 µg/mL) **(C)** Zebrafish treated with mQDs: DOPA (200 µg/mL).

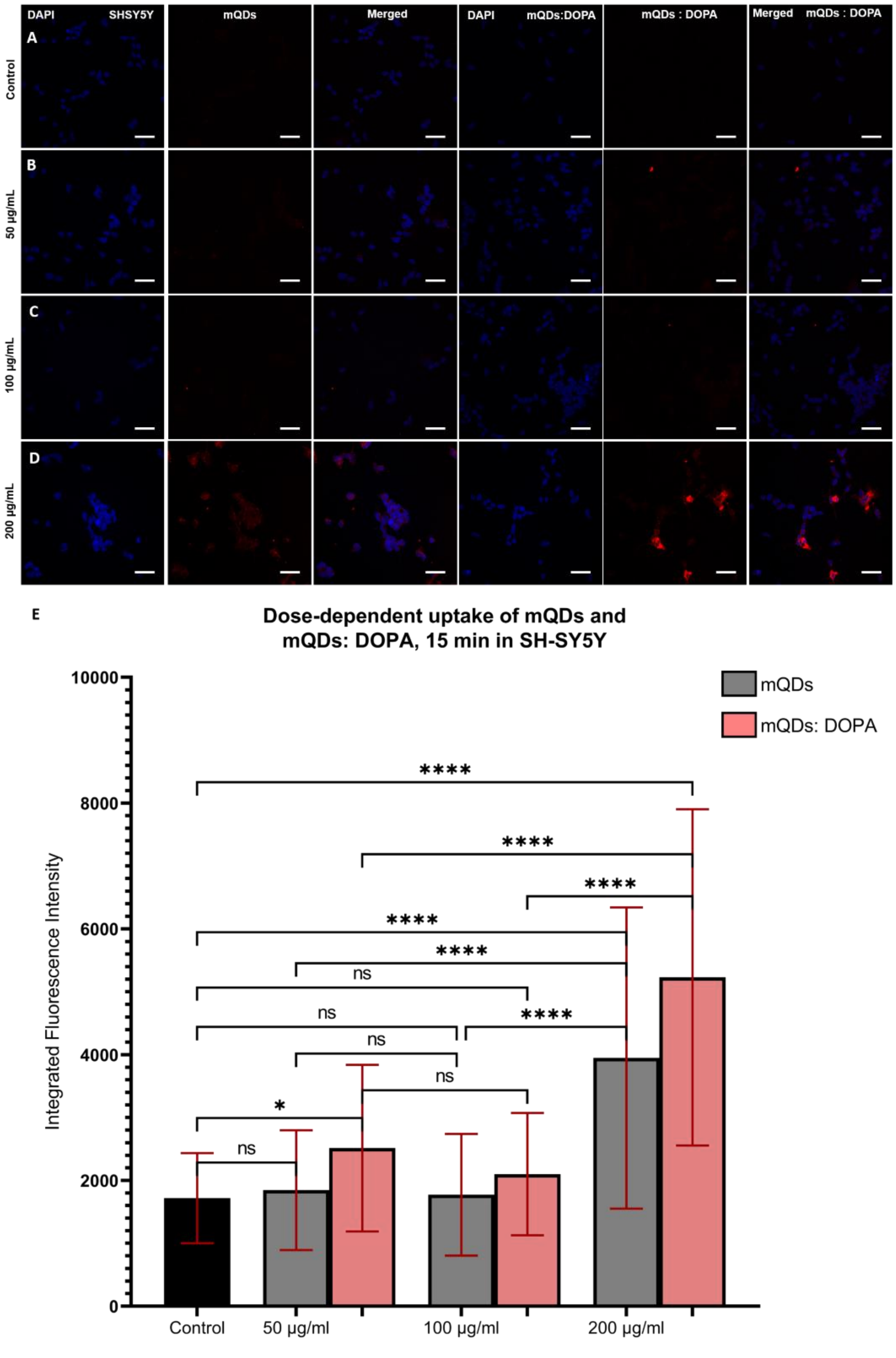
DAPI
SHSY5Y
mQDs
Merged
DAPI
mQDs:DOPA
mQDs : DOPA
Merged
mQDs : DOPA
A
Control
B
50 µg/mL
C
100 µg/mL
D
200 µg/mL
E
Dose-dependent uptake of mQDs and mQDs: DOPA, 15 min in SH-SY5Y
Integrated Fluorescence Intensity
10000
8000
6000
4000
2000
0
mQDs
mQDs: DOPA
****
ns
*
Control
50 µg/ml
100 µg/ml
200 µg/ml

**Figure S10: Dosage-dependent uptake of mQDs and mQDs: DOPA in SHSY-5Y cells. (A)** Imaging of control cells separated into blue (DAPI labelling nucleus) and red (MQDs and mQDs: DOPA fluorescence) channels. **(B)** Imaging of cells with 50 µg/mL dose of mQDs and mQDs: DOPA separated into blue (DAPI labelling nucleus) and red (MQDs and mQDs: DOPA fluorescence) channels. **(C)** Imaging of cells with 100 µg/mL dose separated into blue (DAPI labelling nucleus) and red (MQDs and mQDs: DOPA fluorescence) channels. **(D)** Imaging of cells with 200 µg/mL dose separated into blue (DAPI labelling nucleus) and red (MQDs and mQDs: DOPA fluorescence) channels. **(E)** Quantification of mQDs and mQDs: DOPA uptake depending on dosage applied. Scale bar 5 µm. The following thresholds were used when determining significance. ****: P< 0.0001. *: P< 0.05. ns: P>0.05 and denotes no significance. (n = 50 cells for quantification)

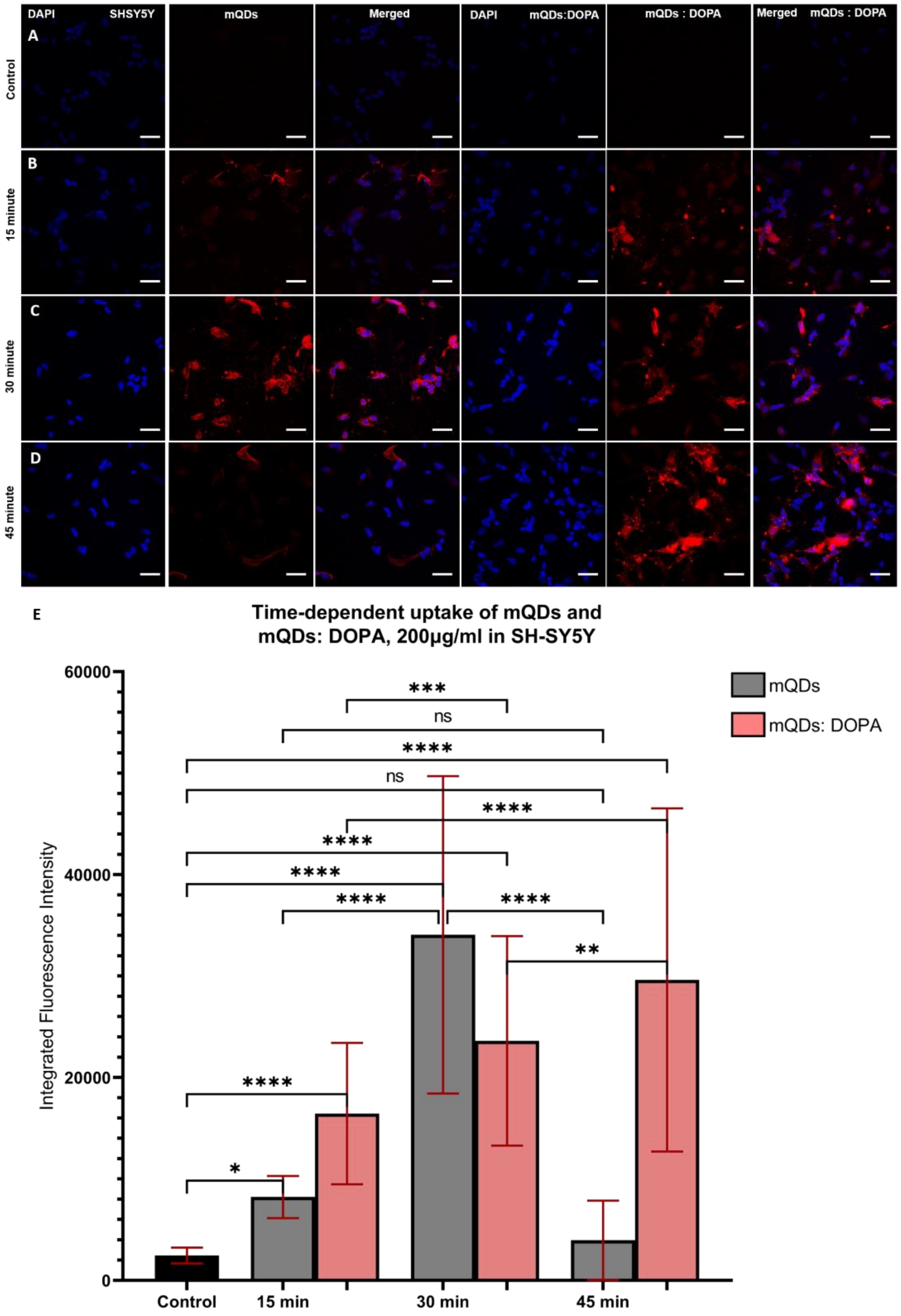
DAPI
SHSY5Y
mQDs
Merged
DAPI
mQDs:DOPA
mQDs : DOPA
Merged
mQDs : DOPA
A
Control
B
15 minute
C
30 minute
D
45 minute
E
Time-dependent uptake of mQDs and
mQDs: DOPA, 200µg/ml in SH-SY5Y
Integrated Fluorescence Intensity
60000
40000
20000
0
Control
15 min
30 min
45 min
mQDs
mQDs: DOPA
***
ns
****
ns
****
****
****
****
****
**
****
*

**Figure S11: Time-dependent uptake of mQDs and mQDs: DOPA in SHSY-5Y cells.**

**(A)** Imaging of control cells separated into blue (DAPI labelling nucleus) and red (MQDs and mQDs: DOPA fluorescence) channels. **(B)** Imaging of cells with 200 µg/mL dose of mQDs and mQDs: DOPA after 15 minutes separated into blue (DAPI labelling nucleus) and red (MQDs and mQDs: DOPA fluorescence) channels. **(C)** Imaging of cells with 200 µg/mL dose after 30 minutes separated into blue (DAPI labelling nucleus) and red (MQDs and mQDs: DOPA fluorescence) channels. **(D)** Imaging of cells with 200 µg/mL dose after 45 minutes separated into blue (DAPI labelling nucleus) and red (MQDs and mQDs: DOPA fluorescence) channels. **(E)** Quantification of mQDs and mQDs: DOPA uptake depending on time. Scale bar 5 µm. The following thresholds were used when determining significance. ****: $P < 0.0001$. ***: $P < 0.001$. **: $P < 0.01$. *: $P < 0.05$. ns: $P > 0.05$ and denotes no significance. (n = 50 cells for quantification)

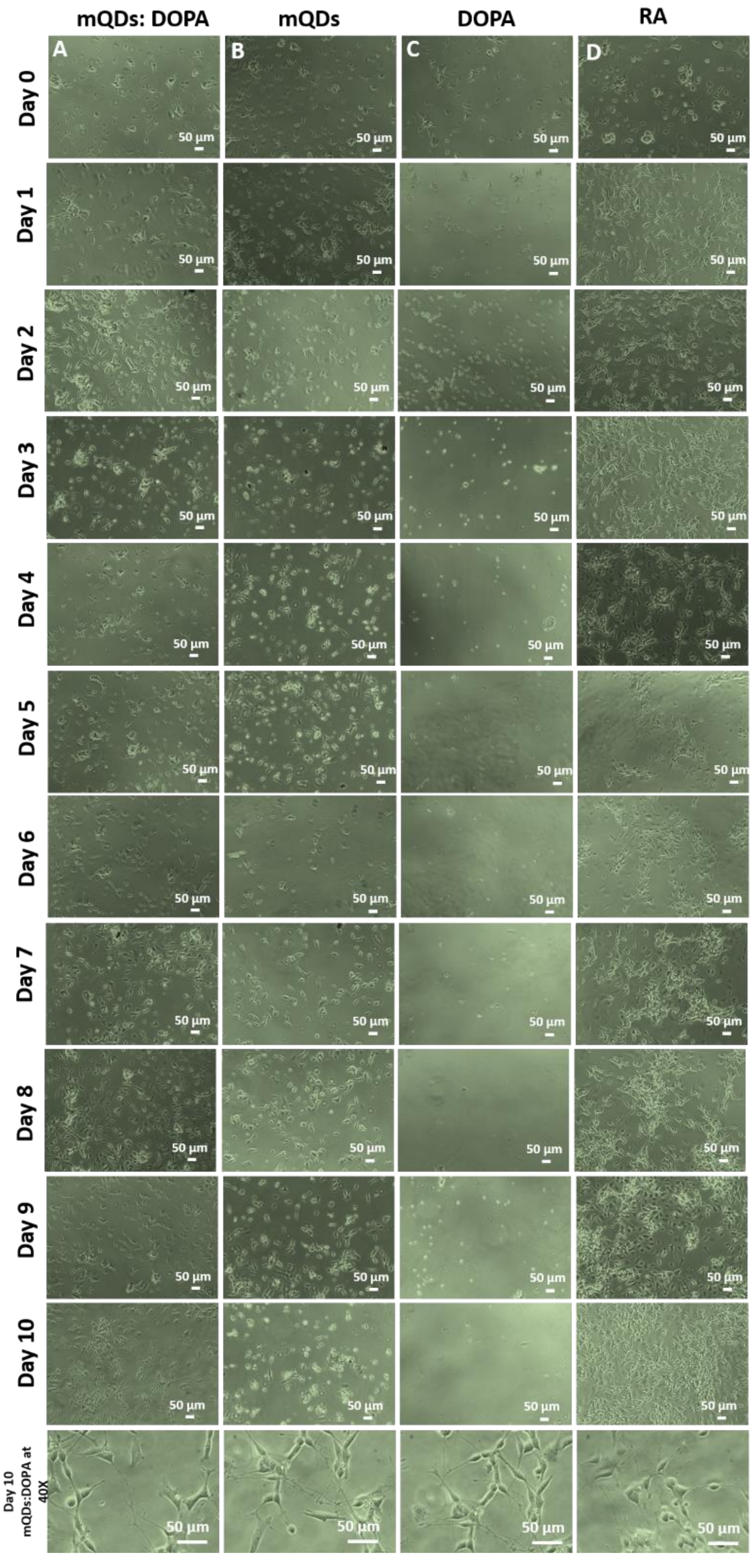

**Figure S12: Day-wise differentiation of SH-SY5Y cells.** Continuous monitoring of cells treated with **(A)** mQDs: DOPA, **(B)** mQDs, **(C)** DOPA, and **(D)** RA (retinoic acid).

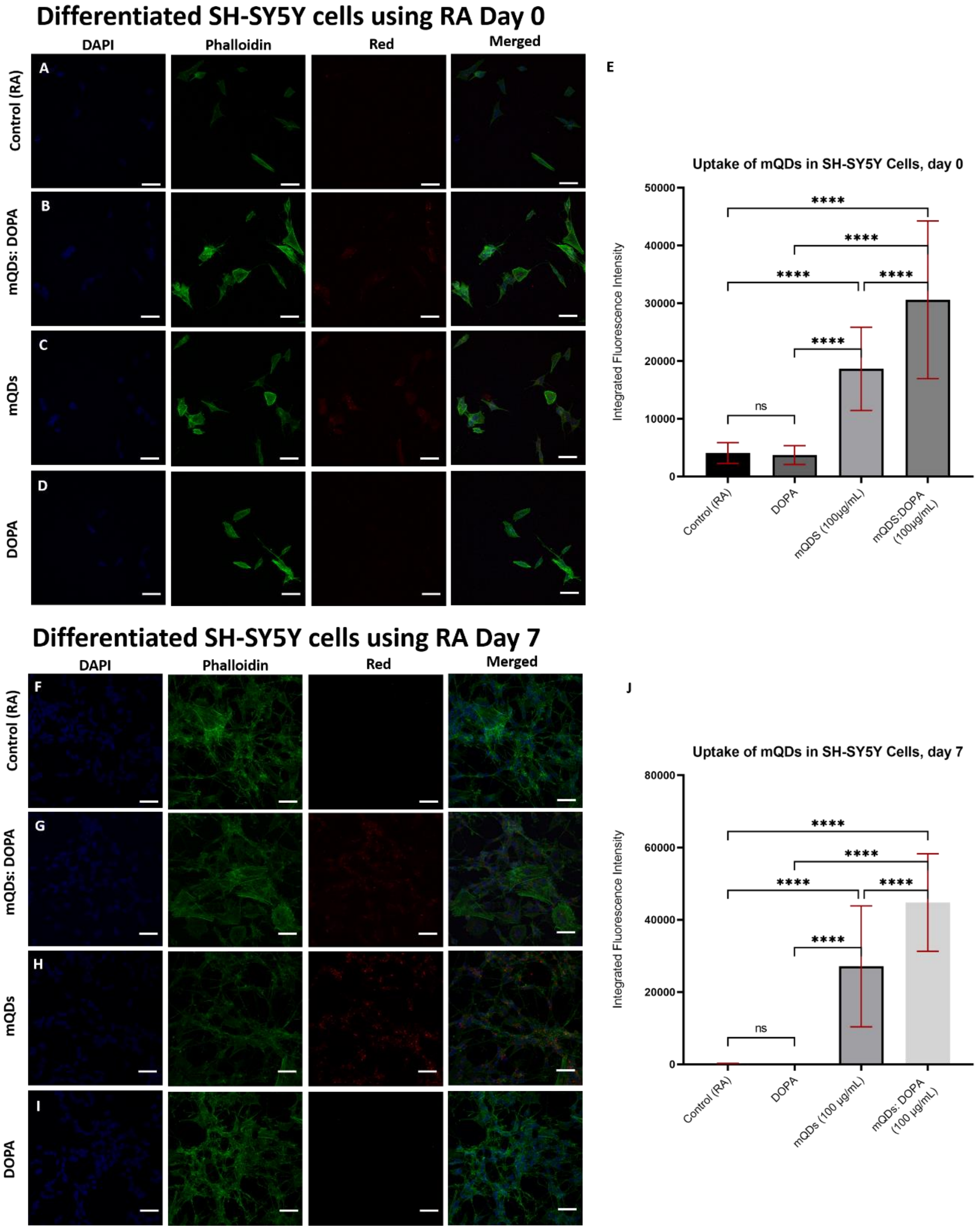

**Figure S13: Differentiated SH-SY5Y cells on day 0 and day 7. (A)** 0-day imaging of control SH-SY5Y cells (retinoic acid) with blue (DAPI labelled nucleus), green (Phalloidin labelled F-actin), and red (mQDs fluorescence) channels. **(B)** 0-day imaging of mQDs: DOPA – treated cells with blue, green, and red channels. **(C)** 0-day imaging of mQDs – treated cells with blue, green, and red channels. **(D)** 0-day imaging of DOPA – treated cells with blue, green, and red channels. **(E)** Quantification of mQDs uptake in SH-SY5Y cells after 0 days. **(F)** 7-day imaging of control SH-SY5Y cells (retinoic acid) with blue (DAPI labelled nucleus), green (Phalloidin labelled F-actin), and red (mQDs fluorescence) channels. **(G)** 7-day imaging of mQDs: DOPA – treated cells with blue, green, and red channels. **(H)** 7-day imaging of mQDs – treated cells with blue, green, and red channels. **(I)** 7-day imaging of DOPA – treated cells with blue, green, and red channels. **(J)** Quantification of mQDs uptake in SH-SY5Y cells after 7 days.

| | **T = 0 hrs.** | **T = 12 hrs.** | **T = 24 hrs.** |
|---|---|---|---|
| **Control** | 342.5287 µm | 272.6293 µm | 249.005 µm |
| **DOPA** | 411.1943 µm | 401.4847 µm | 405.8488 µm |
| **mQDs 100 µg/mL** | 321.5789 µm | 116.5949 µm | 59.66909 µm |
| **mQDs 200 µg/mL** | 347.3294 µm | 136.1205 µm | 88.68328 µm |
| **mQDs: DOPA 100 µg/mL** | 353.8616 µm | 187.663 µm | 143.8335 µm |
| **mQDs: DOPA 200 µg/mL** | 407.6149 µm | 196.1003 µm | 105.2756 µm |

**Table S2:** Average wound length after 0, 12, and 24 hours observed in scratch assay.
